# IL-1β inhibition partially negates the beneficial effects of diet-induced lipid lowering

**DOI:** 10.1101/2023.10.13.562255

**Authors:** Santosh Karnewar, Vaishnavi Karnewar, Rebecca Deaton, Laura S. Shankman, Ernest D. Benavente, Corey M. Williams, Xenia Bradley, Gabriel F. Alencar, Gamze B. Bulut, Sara Kirmani, Richard A. Baylis, Eli R. Zunder, Hester M. den Ruijter, Gerard Pasterkamp, Gary K. Owens

## Abstract

**Background:** Thromboembolic events secondary to rupture or erosion of advanced atherosclerotic lesions are the leading cause of death in the world. The most common and effective means to reduce these major adverse cardiovascular events (MACE), including myocardial infarction (MI) and stroke, is aggressive lipid lowering via a combination of drugs and dietary modifications. However, little is known regarding the effects of reducing dietary lipids on the composition and stability of advanced atherosclerotic lesions, the mechanisms that regulate these processes, and what therapeutic approaches might augment the benefits of lipid lowering.

**Methods:** Smooth muscle cell (SMC)*-*lineage tracing *Apoe^−/−^* mice were fed a Western diet (WD) for 18 weeks and then switched to a low-fat chow diet for 12 weeks. We assessed lesion size and remodeling indices, as well as the cellular composition of aortic and brachiocephalic artery (BCA) lesions, indices of plaque stability, overall plaque burden, and phenotypic transitions of SMC, and other lesion cells by SMC-lineage tracing combined with scRNA-seq, CyTOF, and immunostaining plus high resolution confocal microscopic z-stack analysis. In addition, to determine if treatment with a potent inhibitor of inflammation could augment the benefits of chow diet-induced reductions in LDL-cholesterol, SMC*-*lineage tracing *Apoe^−/−^* mice were fed a WD for 18 weeks and then chow diet for 12 weeks prior to treating them with an IL-1β or control antibody (Ab) for 8-weeks.

**Results:** Lipid-lowering by switching *Apoe^−/−^* mice from a WD to a chow diet reduced LDL-cholesterol levels by 70% and resulted in multiple beneficial effects including reduced overall aortic plaque burden as well as reduced intraplaque hemorrhage and necrotic core area. However, contrary to expectations, IL-1β Ab treatment resulted in multiple detrimental changes including increased plaque burden, BCA lesion size, as well as increased cholesterol crystal accumulation, intra-plaque hemorrhage, necrotic core area, and senescence as compared to IgG control Ab treated mice. Furthermore, IL-1β Ab treatment upregulated neutrophil degranulation pathways but down-regulated SMC extracellular matrix pathways likely important for the protective fibrous cap.

**Conclusions:** Taken together, IL-1β appears to be required for chow diet-induced reductions in plaque burden and increases in multiple indices of plaque stability.

**Clinical Perspective:** Although one must be cautious in extrapolating results of mouse studies to humans, the current and our previous studies (Gomez et al. 2018 *Nature Medicine*) suggest that efforts to identify anti-inflammatory therapies for treating patients with advanced atherosclerosis should consider the possibility that inhibiting a given cytokine may have a mixture of beneficial and detrimental effects that vary between individuals.

**What is New?:** In a mouse model of advanced atherosclerosis followed by diet-induced reductions in cholesterol, IL-1β inhibition unexpectedly had multiple detrimental effects including increasing overall plaque burden and decreasing various indices of plaque stability, as well as markedly increasing the number of senescent cells, and cholesterol crystal accumulation in lesions as compared to IgG control Ab treated mice.

**What are the clinical Implications?:** Results suggest that some anti-inflammatory therapies may have limited efficacy for treating patients with advanced atherosclerosis because such therapies inhibit not only detrimental pro-inflammatory responses, but also evolutionarily conserved beneficial inflammatory processes, which play a critical role in resistance to pathogenic microorganisms, in tissue repair following injury, and resolution of inflammation. We propose that the latter includes clearance of senescent cells and cholesterol from advanced atherosclerotic lesions induced by dietary-induced lipid lowering.

## Introduction

Lipid deposition in the artery wall as a consequence of increased LDL-cholesterol and dyslipidemia is one of the defining characteristics of atherosclerosis. Atherosclerotic plaques characterized by a thick, extracellular matrix-rich fibrous cap with a higher ratio of ACTA2^+^ cells relative to CD68^+^ cells, presumed to be smooth muscle cells (SMC), and macrophages (MΦ) respectively, are more stable^1,2^. A recent study by our lab showed that in murine and human lesions, 20% to 40% of ACTA2^+^ fibrous cap cells, respectively, are derived from non-SMC sources, including EC or macrophages that have undergone an endothelial-to-mesenchymal transition (EndoMT) or a macrophage-to-mesenchymal transition (MMT)^3^. Given that rupture of unstable plaques is the main pathology underlying myocardial infarction (MI) and stroke^4^, it is no surprise that lipid lowering therapies including dietary modifications and statins are widely prescribed. However, despite aggressive lipid lowering, there remain large subsets of patients at increased risk for a MACE^5,6^. Given the compelling evidence that inflammation plays a key role in development of atherosclerosis^7,8^ a dominant hypothesis in the field has been that lipid lowering, combined with suppression of inflammation, would be highly effective in treating patients with advanced disease by generating smaller, more stable plaques^9^.

To test this hypothesis, the Canakinumab Anti-inflammatory Thrombosis Outcome Study (CANTOS) trial identified patients with advanced cardiovascular disease and persistent systemic inflammation despite aggressive lipid lowering and treated them with a control antibody or one of three dosages of canakinumab, an interleukin-1β (IL-1β) neutralizing antibody. Canakinumab, has potent anti-inflammatory effects and has been approved for clinical use in rheumatological disorders^10,11^. In the CANTOS trial, all dosages markedly reduced systemic inflammation as indicated by reduced hsCRP in plasma but only the intermediate dosage achieved a significant reduction in the primary endpoint (a composite of non-fatal MI, non-fatal stroke, and CV death). However, there was no benefit in CV death or overall mortality and there was a 40% increase in lethal infections^12^. Interestingly, a subsequent study of the CANTOS data published in *Lancet* reported that patients achieving an on-treatment hsCRP <2 mg/L had a remarkable 25% reduction in the primary cardiovascular endpoint and a 31% reduction in cardiovascular-related and overall mortality^13^. Moreover, canakinumab appears to be even more effective in patients with Tet2 mutations linked to clonal hematopoiesis^14^. Therefore, canakinumab may well be a valuable treatment option for select cohorts of patients who exhibit effective reductions in blood lipids but have persistently elevated inflammatory biomarkers and specific genetic polymorphisms^13^.

Previous studies have shown that long-term global suppression of inflammation with corticosteroids or the cox-2 inhibitor Vioxx is associated with marked increases in the risk for myocardial infarction, stroke, and heart failure^15,16^. Although the mechanism of action of canakinumab, Vioxx, and corticosteroids are quite different, these data highlight our relatively poor understanding of the effects of suppressing inflammation within advanced atherosclerotic lesions. However, one must consider the possibility that these therapies inhibit not only detrimental pro-inflammatory responses, but also evolutionarily conserved beneficial inflammatory processes which play a critical role not only in resistance to pathogenic microorganisms but also in tissue repair following injury, and resolution of inflammation.

Consistent with this hypothesis, our lab has provided evidence that inhibition of IL-1β signaling in models of advanced atherosclerosis in mice results in a mixture of beneficial and detrimental changes^17^. Inactivation of IL-1 signaling by global knockout of the *Il1r1* in *Apoe^−/−^* mice resulted in smaller lesions but multiple changes consistent with reduced plaque stability including reduced plaque SMC content and fibrous cap coverage, reduced plaque collagen content, and increased intraplaque hemorrhage. *IL1r1* KO mice also had impaired outward vessel remodeling, leading to reduced lumen size^18^. More importantly, our lab also found that treatment of SMC lineage tracing *Apoe^−/−^* mice with advanced lesions for 8-weeks with a murine anti-IL-1β antibody resulted in a >50% reduction in SMC number, ACTA2 content, and collagen content within the fibrous cap as well as reduced SMC but increased macrophage proliferation within lesions^17^. SMC-specific KO of *IL1r1* in Western diet (WD) fed *Apoe^−/−^* mice resulted in a beneficial reduction in lesion size (>70%), but the plaques failed to develop a SMC-rich α-actin (ACTA2^+^) protective fibrous cap^17^. Interestingly, EC-specific KO of *IL1r1* in Western diet fed *Apoe^−/−^* mice resulted in increased lesion size indicating that *IL1r1* signaling in EC is beneficial not detrimental as anticipated^3^. Taken together, the preceding results show that IL-1β-dependent pro-inflammatory signaling may have beneficial or detrimental effects on the pathogenesis of atherosclerosis in different cell types or even within a given cell type at different stages of disease progression.

However, these pre-clinical animal model studies were completed in the context of severe hyperlipidemia and inflammasome activation whereas subjects in human clinical trials^12,13,19,20^ typically have reduced lipids due to aggressive statin treatment combined with dietary changes. As such, we sought to re-examine the effects of a murine IL-1β antibody in the context of a mouse model that better recapitulates clinical practice wherein elderly patients with advanced atherosclerotic disease have a period of reduced lipids prior to possible secondary interventions. Indeed, despite decades of research showing the clear beneficial effects of aggressive lipid-lowering on atherosclerosis^21,22^ how lipid reduction impacts established atherosclerotic lesions including the behavior of key cell types that regulate lesion stability is poorly understood. In studies herein, we test the hypothesis that *treatment with an anti-inflammatory IL-1β antibody will augment the beneficial effects of lipid lowering in Apoe^-/-^ mice with advanced lesions.* Contrary to this hypothesis, we found that IL-1β antibody treatment for 8 weeks following 12 weeks of chow diet-induced lipid-lowering resulted in multiple detrimental changes as compared to IgG antibody treated control mice. These changes included: i) increased plaque burden in the aorta compared to 18w WD+12w CD and age and diet matched IgG control mice [Fig.1E]; ii) Increased BCA lesion size as compared to age and time of diet matched IgG control mice [Fig. 1F]; iii) increased accumulation of cholesterol crystals compared to 18w WD+12w CD group and age and time of diet matched IgG control mice [Fig.1G]; and iv) evidence of reduced plaque stability including increased necrotic core area [Fig.1H], increased intra-plaque hemorrhage [Fig.2C], reduced collagen content in the fibrous cap [Fig.2E], reduced ACTA2 cap thickness per lesion area [Fig.3B], reduced SMC cells in the fibrous cap [Fig.3C], decreased transition of SMC to an ECM producing phenotype [Fig.3D], increased Mac-2 (LGALS3+) in the fibrous cap and increased accumulation of senescent cells [Supplemental figure II], as compared to 18w WD+12w CD and age and time of diet matched IgG controls. As such, these studies reveal an unanticipated beneficial role for IL-1β in lesion remodeling and stabilization in response to diet-induced decreases in lipids in mice that is mediated in part by the prevention of cholesterol crystal formation.

**Figure 1.**
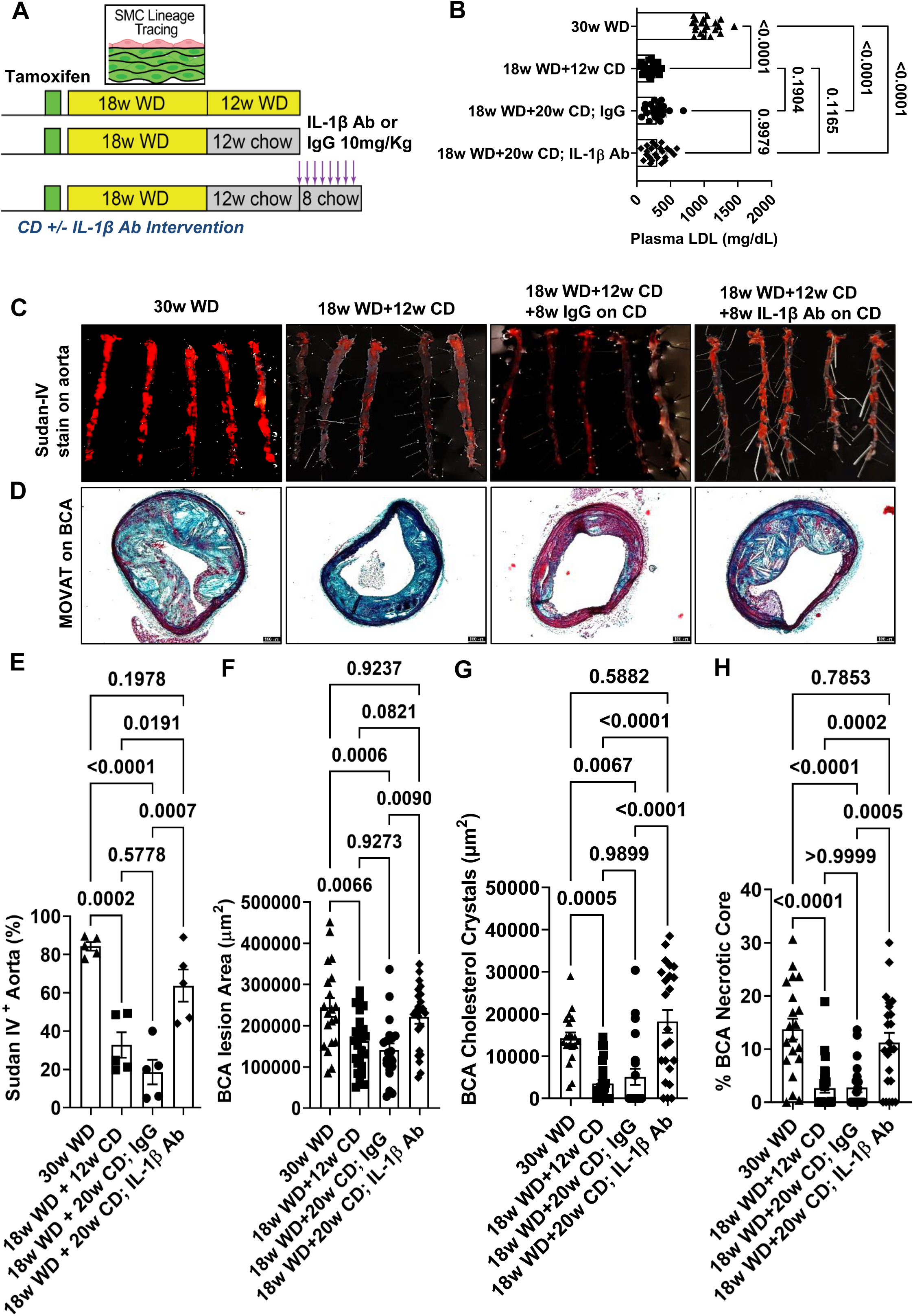
IL-1β Ab treatment of *Apoe^-/-^* mice after chow diet-induced lipid lowering resulted in an increased plaque burden, BCA lesion size, BCA necrotic core area, and BCA cholesterol crystal accumulation, as compared to age and time of diet matched IgG control treated mice. **A**, Experimental design, SMC lineage tracing *Apoe^−/−^* mice were injected with tamoxifen at 6 to 8 weeks of age and subsequently placed on a WD for 18 weeks to induce advanced lesion formation followed by switching mice to chow diet for 12 weeks or one group continued receiving WD for another 12 weeks as 30weeks WD progression group. Chow diet-switched mice were then randomized to being treated with a murine IL-1β neutralizing Ab or an isotype-matched IgG control Ab for 8 weeks while mice continued on chow diet. **B**, Plasma LDL cholesterol. **C**, Sudan-IV staining on aortas. **D**, Representative Movat images. **E**, Sudan-IV+ area quantification. **F**, BCA lesion area **G**, Quantification of cholesterol crystals in the BCA. **H**, Percentage necrotic core area quantified from BCA. Error bars represent mean ± SEM, p-values displayed refer to one-way ANOVA with multiple comparisons. Each triangle (n=20), square (n=26), circle (n=21) and diamond (n=23) shape on the graph indicates data from an individual mouse except for Sudan-IV stain (n=5).

**Figure 2.**
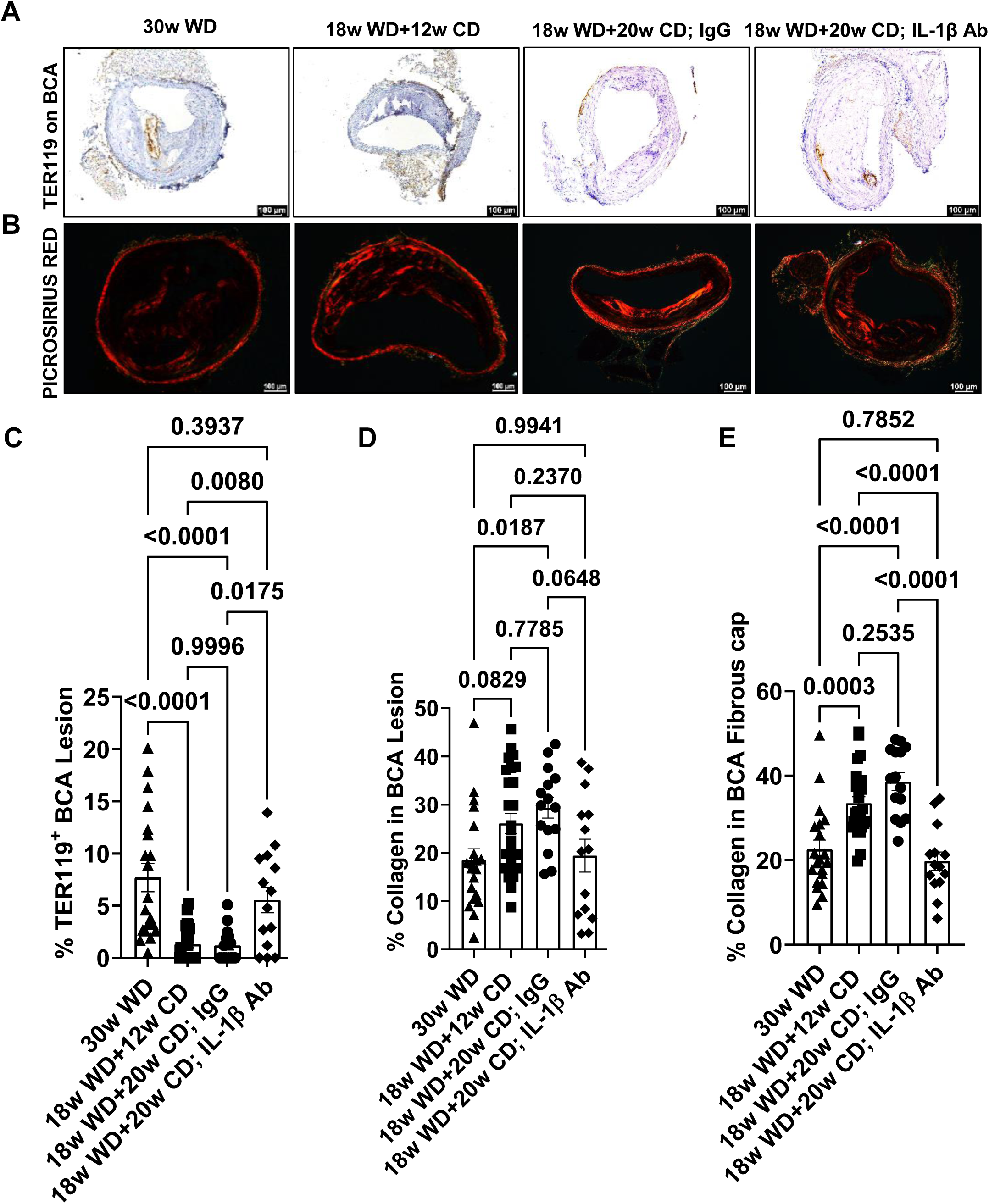
Treatment of *Apoe^−/−^* mice with an IL-1β neutralizing Ab after chow diet-induced lipid lowering resulted in an increased intraplaque hemorrhage (Ter119+) in the BCA lesions, and reduced collagen content in the fibrous cap of BCA. Same experimental groups as shown in Figure 1. **A**, TER119 stained BCA lesions. **B**, Picrosirius red stained BCA lesions. **C**, Percentage Intraplaque hemorrhage (TER119^+^) lesions. **D**, Percentage of collagen in BCA lesion. **E**, Percentage of collagen in BCA fibrous cap. Error bars represent mean ± SEM, p-values displayed refer to one-way ANOVA with multiple comparisons. Each triangle (n=20), square (n=26), circle (n=15) and diamond (n=14) shape on the graph indicates data from an individual mouse.

**Figure 3.**
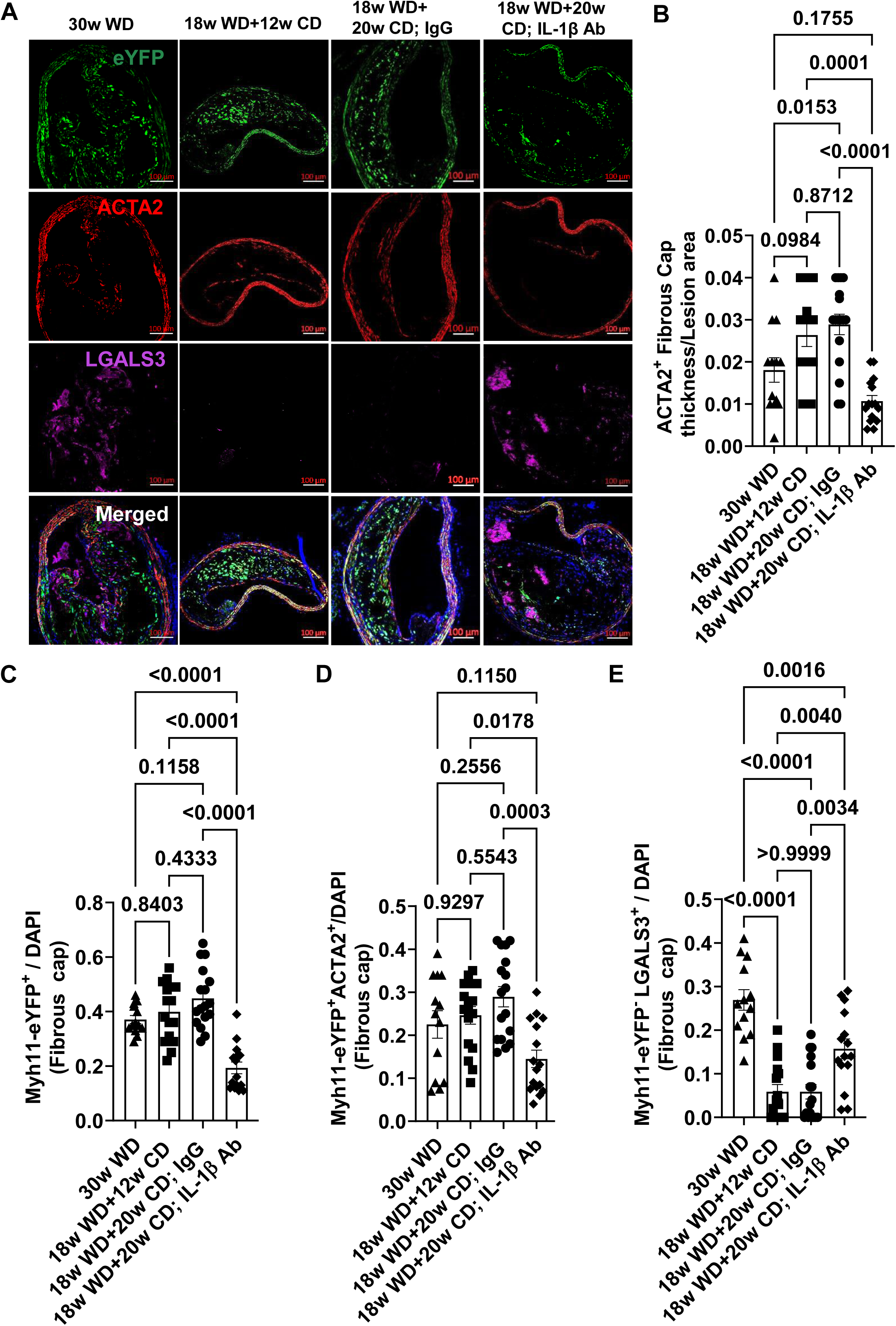
IL-1β Ab treatment of *Apoe^-/-^* mice after chow diet-induced lipid lowering resulted in reduced ACTA2^+^ fibrous cap thickness and reduced SMC and SMC-derived ACTA2^+^ cells but increased LGALS3^+^ cell investment into the fibrous cap of **BCA lesions.** Same experimental groups as shown in Figure 1. **A**, Confocal images of BCA lesions stained with eYFP, ACTA2, LGALS3, and DAPI. **B**, ACTA2^+^ fibrous cap thickness per BCA lesion area. **C**, Myh11-eYFP^+^ (SMC) of all (DAPI^+^) cells in the fibrous cap. **D**, SMC derived ACTA2^+^ cells of all (DAPI^+^) cells in the fibrous cap. **E**, non-SMC derived LGALS3^+^ (Myh11-YFP^-^LGALS3^+^/DAPI) cells of all (DAPI) cells in the fibrous cap. Error bars represent mean ± SEM, p-values displayed refer to one-way ANOVA with multiple comparisons. Each triangle (n=13), square (n=16), circle (n=17) and diamond (n=15) shape on the graph indicates data from an individual mouse.

## Methods

### Animals, diet, and treatment

The University of Virginia Animal Care and Use Committee approved animal protocols. The *Myh11-CreERT2 R26R-eYFP Apoe^−/−^* mice used in the present study have been described in previous studies^23,17,24^. In *Myh11-CreERT2 R26R-eYFP Apoe^−/−^* mice cre recombinase was activated with a series of ten tamoxifen injections (1mg/day/mouse; Sigma Aldrich, T-5648) over a 2-weeks period. One week after the tamoxifen treatment, mice were switched to a high fat western diet (WD), containing 21% milk fat and 0.2% cholesterol (Harlan Teklad; TD.88137; 17.3% protein, and 21.2 % fat, 0.2% Cholesterol) for either 30 weeks in a progression model or 18 weeks of WD followed by 12 weeks of standard chow diet-induced lipid lowering (Harlan Teklad TD.7012; 19.1% protein, and 5.8 % fat, and 0% cholesterol) model. For our intervention study, we fed Myh*11-Cre ERT2 R26R-eYFP Apoe^−/−^* mice a WD for 18 weeks followed by a standard chow diet for 12 weeks, and mice were randomized into two groups and then treated with a murine IL-1β (Provided by Novartis), or isotype-matched IgG control antibody [administered intraperitoneally (I.P.) at 10mg/kg] for 8 more weeks on chow diet.

### Detailed methods were included in supplemental methods

#### Statistical analysis

Statistics were performed using GraphPad Prism 9. Data normality was determined using the Kolmogorov-Smirnov test. For comparison of two groups of continuous variables with normal distribution and equal variances, two-tailed unpaired Student’s t-tests were performed with a confidence level of 95%. For comparison of two groups of continuous variables with normal distribution and unequal variances, two-tailed unpaired Student’s t-tests followed by a Welch’s correction were performed with a confidence level of 95%. Two-tailed unpaired Mann-Whitney U-tests with a confidence level of 95% were conducted if data were non-normally distributed. For multiple group comparison, we performed one-way ANOVA with Tukey’s multiple comparisons. For categorical data (intraplaque hemorrhage analysis), we used fisher’s exact test. The number of mice used for each analysis is indicated in the figure legends and on the graphs. Statistical significance threshold was set at P≤0.05. All data are presented as the mean ± SEM.

## Results

### IL-1β inhibition after chow diet-induced lipid lowering resulted in increased plaque burden, senescence, cholesterol crystal accumulation, and necrotic core area

Our initial goal was to determine if switching mice with advanced atherosclerosis to a low-fat chow diet is associated with beneficial changes including reductions in lesion size and plaque burden as well as increases in indices of plaque stability. We sought to establish a mouse model that better replicates human clinical scenarios wherein hyperlipidemic patients with advanced atherosclerosis are initially put on lipid lowering therapies and diets prior to any secondary interventions. We accomplished this by feeding SMC lineage-tracing *Apoe^−/−^* mice a WD for 18 weeks which we have previously shown induces formation of advanced atherosclerotic lesions in the BCA and aortic tree^17,3^. We then switched mice to a standard chow diet for 12 weeks and compared them to littermate control mice fed a WD for 18 or 30 weeks (Supplemental Figure I), the latter being the longest time period allowed by our Animal Care and Use Committee. Mice fed a WD for 18 weeks followed by 12 weeks of chow diet had a mean plasma LDL concentration of 213 mg/dL as compared to 1034 mg/dL for age matched littermate controls on a 30w WD or 1030 mg/dL on an 18w WD (Supplemental Figure I). We found similar results with plasma cholesterol and triglyceride levels (Supplemental Figure I). However, we did not see any change in HDL cholesterol levels between the groups (Supplemental Figure I). Consistent with human studies showing little or no decrease in lesion size with aggressive statin treatment^25,26,27^, we found no change in brachiocephalic artery (BCA) lesion area in mice fed a WD for 18 weeks followed by 12 weeks of chow diet (referred to as the “18w WD + 12w CD”) mice as compared to littermate control mice fed a WD for 18 weeks (Supplemental Figure I). However, we saw decreased necrotic core area and intraplaque hemorrhage in the 18w WD+12w CD group compared to the 18w WD group (Supplemental Figure I).

We previously showed that treatment of mice with advanced atherosclerotic lesions with an anti-IL-1β neutralizing Ab resulted in lesions that had multiple features consistent with reduced stability including a marked decrease in collagen content and SMC number within the fibrous cap^17^. However, our previous study involved treating mice with severe hyperlipidemia and thus may not apply to human clinical scenarios or clinical trials wherein the first line of treatment of patients with advanced atherosclerotic disease is aggressive lipid lowering through use of statins and dietary changes. Thus, we hypothesized that the multiple detrimental effects of IL-1β Ab treatment on advanced lesions of WD-fed *Apoe^−/−^* mice would be abrogated or even reversed after long-term reductions in blood lipid levels.

We tested this hypothesis using our chow diet-switch model. *Myh11-eYFP^+^ Apoe^−/−^* mice were fed a WD for 18 weeks to induce advanced lesion formation followed by switching mice to chow diet for 12 weeks. Mice that continued on a WD for 30 weeks served as 30w WD progression group. 18w WD+12w chow diet-switched mice were then randomized to being treated with a murine IL-1β neutralizing Ab or an isotype-matched IgG control Ab for 8 weeks while mice continued on a chow diet (Figure 1A). Chow diet-switch reduced plasma LDL levels compared to the 30w WD progression group, however IL-1β Ab did not change the LDL cholesterol levels compared to IgG control Ab (Figure 1B). The plaque burden measured with Sudan-IV was markedly reduced in 18w WD +12w CD group mice as compared to age matched control mice fed a WD for 30 weeks (30w WD “progression group”) consistent with overall lesion regression (Figure 1C and 1E). Surprisingly, although control Ab-treated mice (18w WD+20w CD IgG) showed the expected decrease in plaque burden as shown in Figure 1C and 1E, the mice treated with IL-1β Ab did not (Figure 1C and 1E). More surprisingly, the mice treated with IL-1β Ab showed a marked increase in plaque burden as compared to 18w WD+12w CD group (Figure 1C and 1E). Similarly, 18w WD+12w CD and 18w WD+20w CD IgG groups had evidence of less cell senescence based on reduced aortic SAβG^+^ staining as compared to 30w WD progression group. However, this chow diet induced decrease was not seen in 18w WD+20w CD mice treated with IL-1β Ab (Supplemental Figure II). In addition, the BCA lesion area of the 18w WD+12w CD group mice and 18w WD+20w CD IgG group mice were markedly decreased as compared to mice fed a WD for 30 weeks. The mice treated with IL-1β Ab also did not have reduced lesion area (Figure 1D, 1F and Supplemental Figure III-Figure VII). Furthermore, cholesterol crystal accumulation and the necrotic core area were markedly decreased in 18w WD+12w CD group mice as compared to 30w WD “progression group” (Figure 1G, 1H and Supplemental Figure III-VII). In addition, although control Ab-treated mice (18w WD+20w CD IgG) had reduced cholesterol crystal and necrotic core areas within BCA lesions as shown in Figure 1G and 1H, mice treated with IL-1β Ab did not (Figure 1G, 1H and Supplemental Figure III-VII). Indeed, the mice treated with IL-1β Ab showed a marked increase in BCA cholesterol crystal and BCA necrotic core areas as compared to the 18w WD+12w CD group (Figure 1G, 1H and Supplemental Figure III-VII). Taken together studies indicate that IL-1β Ab treatment negated multiple beneficial effects of a lipid lowering diet.

### IL-1β Ab treatment of *Apoe^-/-^* mice after chow diet-induced lipid lowering resulted in reduced indices of plaque stability

We also observed multiple beneficial changes in indices of plaque stability in the 18w WD +12w CD and 18w WD+20w CD IgG groups versus the 30w WD progression and 18w WD+20w CD IL-1β Ab treated mice. These beneficial changes included reduced intra-plaque hemorrhage, and increased collagen content in the fibrous cap (Figure 2A-2E). However, IL-1β Ab treatment negated these beneficial changes (Figure 2A-2E). To ascertain if the preceding changes were also associated with changes in the cell types in the fibrous cap, we stained the BCA lesions of 30w WD and 18w WD+12w CD, 18w WD+20w CD IgG, and 18w WD+20w CD IL-1β Ab for eYFP, ACTA2, LGALS3, and DAPI (Figure 3A). Notably, ACTA2^+^ fibrous cap thickness was reduced with IL-1β Ab but not with control-Ab treatment (Figure 3B). Furthermore, SMC (Myh11-eYFP^+^/DAPI), SMC– derived ACTA2^+^ (Myh11-eYFP^+^ ACTA2^+^/DAPI) cells, were not different in the fibrous cap of 18w WD+12w CD group compared to 30w WD progression group. However, SMC and SMC–derived ACTA2^+^ cells were reduced in the fibrous cap of mice treated with IL-1β Ab compared to control-Ab treatment or 18w WD+12w CD groups (Figure 3C-3D). Non SMC-derived Lgals3^+^ cells (Myh11-eYFP^+^LGALS3^+^/DAPI) presumed to be macrophages^3^ were reduced in the fibrous cap of 18w WD+12w CD group compared to 30w WD progression group (Figure 3E). Interestingly, IL-1β Ab treatment resulted in increased LGALS3+ cells in the fibrous cap compared to IgG control group mice or 18w WD+12w CD group mice (Figure 3E). Taken together, results provide evidence that IL-1β plays a critical role in promoting improvements in indices of plaque stability during chow diet-induced lipid lowering.

### IL-1β inhibition resulted in an increased fraction of osteochondrocyte marker+ cells in BCA lesions

To determine potential mechanisms for the beneficial changes in indices of plaque stability with chow diet-induced lipid lowering, we analyzed changes in the cellular composition of advanced lesions by Cy-TOF analysis of the BCA region from 30w WD and 18w WD+12w CD SMC-lineage tracing *Apoe^-/-^* mice (Supplemental Figure VIIIA). We used a 26 marker antibody panel previously described^24^ [see supplemental methods section and supplemental Figure VIIIB]. We found 11 clusters including immune cells, SMC, and EC marker+ clusters (Supplemental Figure VIIIC). Interestingly, cluster-2 (CD45^+^ CD86^-^) presumed to be T-cells was increased in 18w WD+12w CD group compared to 30w WD group (Supplemental Figure VIIID). Importantly, the cluster-5 which expresses multiple macrophage markers including F4/80^+^, CD11b^+^, CX3CR1^+^, CD45^+^, CD86^+^, CD124^+^, CD184^+^, CD206^+^, CD140b^+^, and RUNX2^+^, was decreased in the 18w WD+12w CD group compared to 30w WD group (Supplemental Figure VIIIE). In addition, cluster-9, which includes CD45+ and CD11b+ macrophages was also decreased in 18w WD+12w CD group compared to 30w WD group (Supplemental Figure VIIIF). Furthermore, cells in cluster-8, which express inflammatory and osteochondrocyte markers including IL-6^+^, CD44^+^, CD140a^+^, CD140b^+^, RUNX2^+^, IFN-g^+^, TRPV4^+^, OCT3/4^+^, and LY6A^+^ was decreased in 18w WD+12w CD fed mice as compared to 30w WD-fed mice (Supplemental Figure VIIIG). Taken together, results show that the 18w WD+12w CD group had reduced numbers of macrophages and osteochondrocytes within BCA lesions compared to the age-matched 30w WD progression group.

We also performed the Cy-TOF analysis on BCA region samples from IL-1β Ab versus IgG control Ab treated mice (Figure 4A). We used the same 26 marker antibody panel as shown in previous section (Supplemental Figure VIII). A heat map showing the results of these Cy-TOF analyses is shown in Figure 4B. We found 10 clusters including immune cells, and SMC marker+ clusters (Figure 4C). Interestingly, cluster-2 (CD45^+^ CD86^-^) which is a T-cell cluster was decreased with IL-1β Ab treatment compared to control-Ab treatment (Figure 4D). In contrast, cluster-1 which express KLF4^+^, LY6A^+^, CD206^+^, CD34^+^, and CD140A^+^ was increased in the IL-1β Ab treatment group compared to the IgG group (Figure 4E). In addition, cluster-3, that includes SMC-derived cells which are GFP+ (SMC), CD200^+^, IL-6^+^, NG2^+^, S100B^+^, IFN-g^+^ and desmin^+^ was increased compared to IgG group (Figure 4F). Furthermore, cluster-5 that also includes SMC-derived (GFP+) cells which express multiple osteochondrocyte markers including TRPV4^+^, S100B^+^, RUNX2^+^, OCT3/4^+^, KLF4^+^, CD200^+^, CX3CR1^+^, IFN-g^+^, NG2^+^, DESMIN^+^, CD184^+^, CD34^+^, and CD117^+^was increased with IL-1β Ab treatment compared to the control-Ab group (Figure 4G).

**Figure 4.**
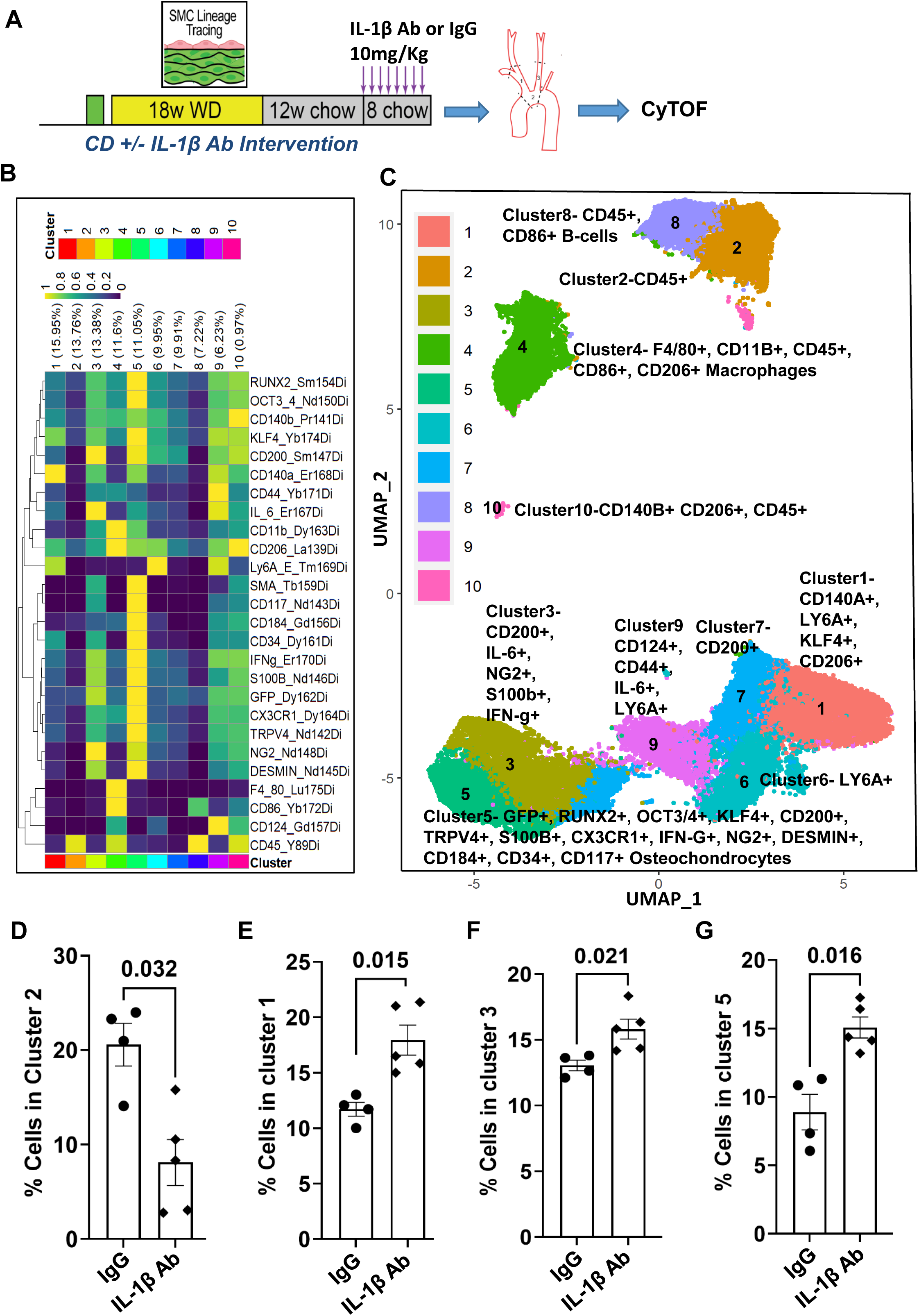
Treatment of *Apoe^−/−^* mice with an IL-1β neutralizing Ab after chow diet-switch induced lipid lowering increased SMC-derived osteochondrocyte marker+ cells. **A**, Experimental design, SMC lineage tracing *Apoe^−/−^* mice were injected with tamoxifen at 6 to 8 weeks of age and subsequently placed on a WD for 18 weeks to induce advanced lesion formation followed by switching mice to chow diet for 12 weeks. Chow diet-switched mice were then randomized to being treated with a murine IL-1β neutralizing Ab or an isotype-matched IgG control Ab for 8 weeks while mice continued on chow diet. **B**, 26-antibody panel Cy-TOF staining on BCA region lesion cells. **C**, BCA region lesion cells are shown as a UMAP colored by cluster from Cy-TOF analysis. **D**, percentage of cells in cluster 2 (CD45+ but negative for the B-cell marker CD86). **E**, percentage of cells in cluster 1 (KLF4+, CD140A+, LY6A+, CD34+, CD206+) **F**, percentage of cells in cluster 3 (GFP+, CD200+, IL-6+, NG2+, S100B+, IFN-g+). **G,** percentage of cells in cluster 5 (GFP+ and osteochondrocyte markers+ cells). Error bars represent mean ± SEM; p-values displayed refer to Mann-Whitney U-test between IL-1β Ab (n=5) and control Ab (n=4) treated groups. Each circle or diamond shape on the graph indicates data from an individual mouse.

### scRNAseq analyses showed that IL-1β Ab treatment of *Apoe^-/-^* mice after chow diet-induced lipid lowering resulted in a reduced fraction of SMC in lesions exhibiting a SMC marker+ phenotype but an increased fraction of the osteochondrocyte marker+ cells

To better understand the mechanisms by which IL-1β inhibition induced multiple detrimental changes in BCA lesions after chow diet-induced lipid lowering compared to IgG controls, we performed scRNA-seq on micro-dissected BCA lesions from *Myh11-eYFP^+^ Apoe^−/−^*mice treated with IL-1β Ab or IgG (Supplemental Figure IX). As previously observed by our lab^24^ and others^28^, we found lesion transcriptomic clusters corresponding to different phenotypic states of SMC, EC, macrophages, and T-cells (Figure 5 and Supplemental Figure IX). Importantly, IL-1β Ab treated mice had a reduced percentage of SMC marker+ cells in cluster-2 which are believed to represent fibrous cap cells that have re-differentiated^24^ (Figure 5C-5E, and Supplemental Figure IX-XI). The IL-1β Ab group had increased proportion of cells in cluster-4 which exhibit an osteochondrocyte phenotype likely to contribute to plaque calcification and to be detrimental for plaque stability (Figure 5D). In addition, the IL-1β Ab treatment group showed higher plaque Von kossa+ calcification compared to IgG control group (Supplemental Figure XII).

**Figure 5.**
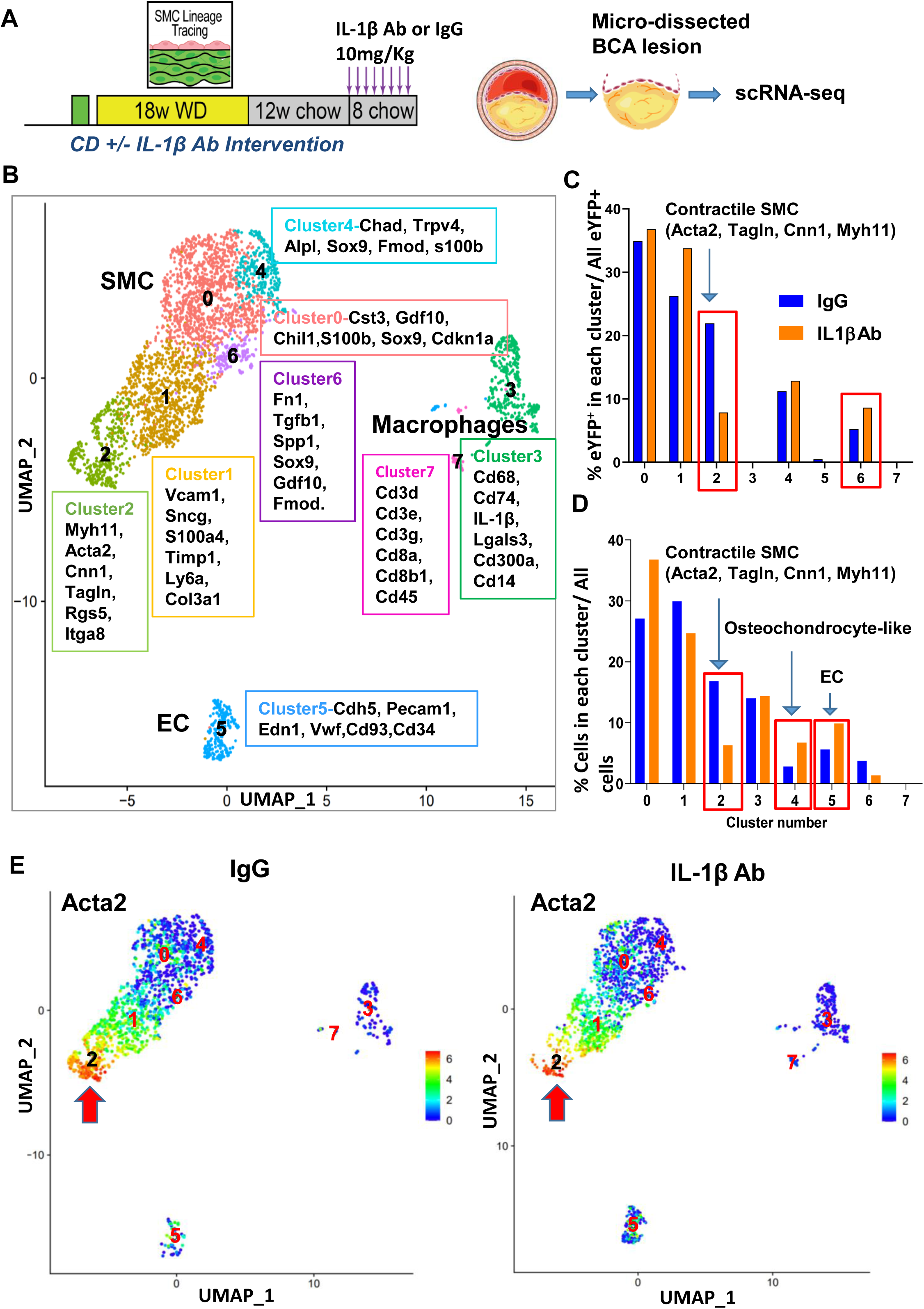
Treatment of *Apoe^−/−^* mice with an IL-1β neutralizing Ab after chow diet-induced lipid lowering resulted in a reduced fraction of SMC marker^+^ lesion cells but an increased fraction of osteochondrocyte marker^+^ cells. **A**, Experimental design, SMC lineage tracing *Apoe^−/−^* mice were injected with tamoxifen at 6 to 8 weeks of age and subsequently placed on a WD for 18 weeks to induce advanced lesion formation followed by switching mice to chow diet for 12 weeks. Mice were then randomized to being treated with a murine IL-1β neutralizing Ab or an isotype-matched IgG control Ab for 8 weeks while mice continued on chow diet. **B**, Cells were harvested for scRNA-seq from micro-dissected advanced BCA lesions and results are shown as a UMAP colored by cluster. **C**, Percentage of SMC (Myh11-eYFP+) lesion cells in each UMAP cluster from each respective group described in B. **D**, Percentage of all lesion cells in each UMAP cluster from each respective group described in B. **E**, Acta2 UMAP analysis shows cells from IgG on left and from IL-1β on the right, and red arrows indicating Acta2^+^ cells were changed in cluster-2.

Given inherent limitations in scRNA-seq analyses related to modest read depth, possible sample biases in isolating single cells from tissues, extremely high cost, and it requiring pooling lesions from 4-6 mice for each experimental and control sample and thus a lack of independent replicates (n=1), we also performed bulk-RNA-seq on micro-dissected BCA lesions from IL-1β Ab (n=4) or IgG control (n=4) treated *Myh11-eYFP+ Apoe^−/−^* mice after chow diet-switch (Figure 6A). Interestingly, reactome pathway analysis showed that inhibition of IL-1β resulted in up-regulation of neutrophil degranulation pathways (Supplemental Figure XIII). In addition, IL-1β inhibition was associated with down-regulation of cell cycle, and extracellular matrix organization pathways (Supplemental Figure XIII). Overall results of differentially expressed genes (DEG) analysis showed that the IL-1β Ab treatment group had upregulated 746 genes and down regulated 725 genes (Figure 6B). The top upregulated genes with IL-1β Ab treatment group were osteochondrocyte and pro-inflammatory SMC markers which is consistent with our SMC-lineage tracing scRNA-seq data (Figures 5B-5C and Supplemental Figure X and XI) from the previous section. Notably, the top up-regulated genes from bulk RNA-seq included *Gdf15 and Klf4* which were also found in cluster2 in our scRNAseq analyses (Figures 5B, Figure 6C and 6D). Additional upregulated genes and their corresponding scRNAseq cluster included *Ly6c1* and *Ly6a* in cluster1 and cluster5 (Figures 5B, Figure 6E and 6F)*, and Ly6e* in cluster1 (Figures 5B and Figure 6G). SNPs near *Klf4* has been identified as coronary artery disease GWAS variant^29^. The top down-regulated genes and their corresponding scRNAseq cluster include *Itgb8* and *Fgf2* (Figure 6H-6I) which was found in cluster2 which are likely to be SMC-derived fibrous cap cells. Taken together, these studies indicate that IL-1β Ab treatment of *Apoe^-/-^* mice after chow-diet induced lipid lowering resulted in a reduced fraction of SMC within lesions exhibiting a SMC marker+ plaque-stabilizing phenotype and an increased fraction of SMC that have transitioned to an osteochondrocytic phenotype likely to be detrimental for lesion pathogenesis.

**Figure 6.**
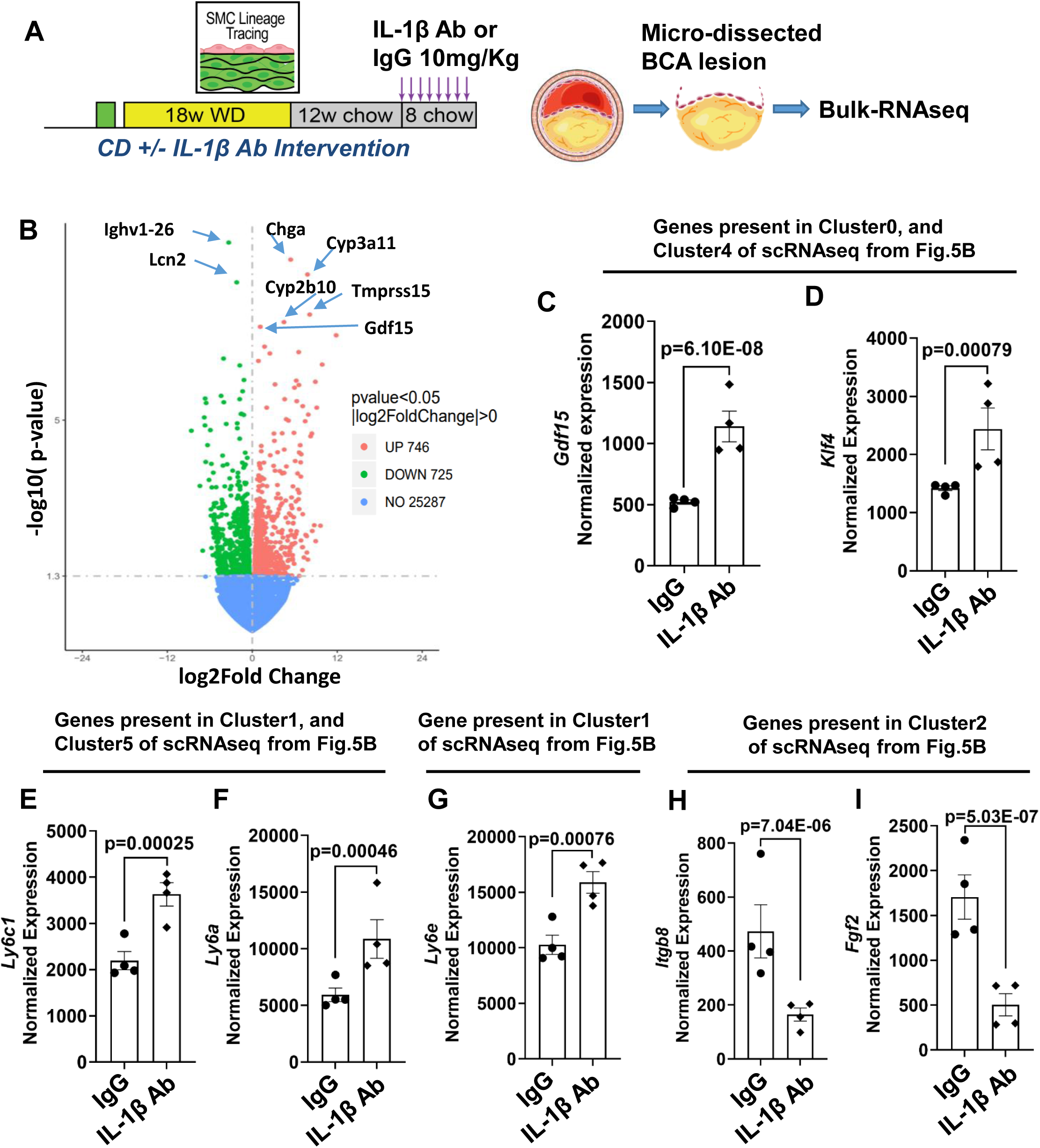
Treatment of *Apoe^−/−^* mice with an IL-1β neutralizing Ab after chow diet-induced lipid lowering resulted in decreased expression of SMC marker genes and Increased expression of osteochondrocyte marker genes in BCA lesions. **A**, Experimental design, SMC lineage tracing *Apoe^−/−^* mice were injected with tamoxifen at 6 to 8 weeks of age and subsequently placed on a WD for 18 weeks to induce advanced lesion formation followed by switching mice to chow diet for 12 weeks. Mice were then randomized to being treated with a murine IL-1β neutralizing Ab (n=4) or an isotype-matched IgG control Ab (n=4) for 8 weeks while mice continued on chow diet. Bulk-RNA-seq was then performed on micro dissected advanced BCA lesions. **B**, Volcano plot of DEG up and down regulated genes. C*, Gdf15*. D, *Klf4*. E, *Ly6c1*. **F**, *Ly6a*. **G**, *Ly6e*. **H**, *Itgb8*. **I**, *Fgf2*. **J**, Heat map of GSEA analysis showing 50 up and 50 down regulated genes with IL-1β neutralizing Ab versus IgG. Error bars represent mean ± SEM; p-values displayed refer to Mann-Whitney U-test between IL-1β Ab (n=4) and control Ab (n=4) treated groups. Each circle or diamond shape on the graph indicates data from an individual mouse.

**Figure 7.**
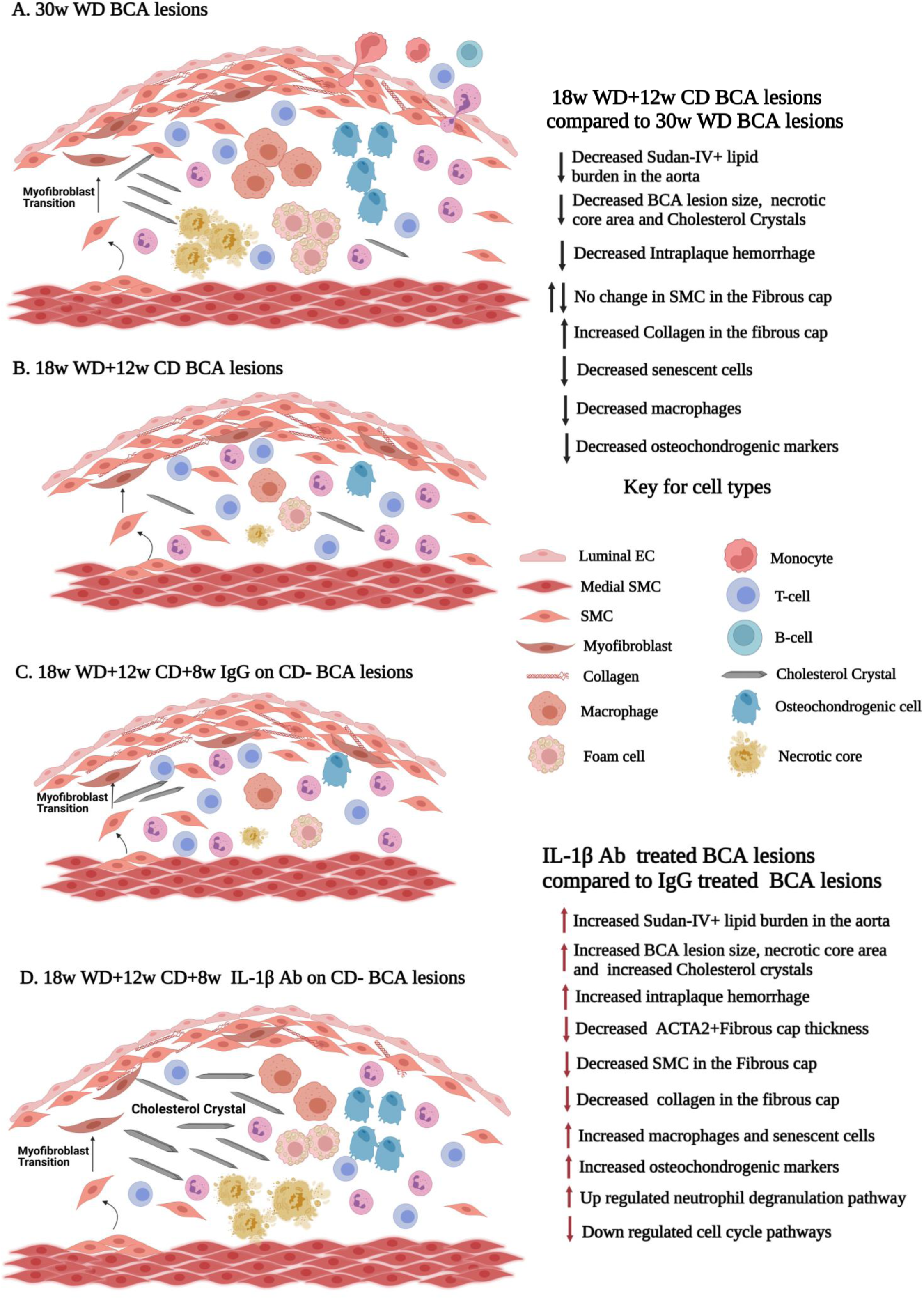
Graphical Images showing treatment of *Apoe^−/−^* mice with an IL-1β neutralizing Ab after chow diet-induced lipid lowering resulted in multiple detrimental effects on indices of plaque stability. **A**, shows SMC lineage tracing *Apoe^−/−^* mice were injected with tamoxifen at 6 to 8 weeks of age and subsequently placed on a WD for 30 weeks (30w WD controls). **B**, shows after tamoxifen at 6 to 8 weeks of age mice were placed on a Western diet (WD) for 18 weeks and then switching mice to chow diet for 12 weeks (18w WD+12w CD) reduced lesion size, reduced necrotic core area, and reduced macrophages compared to 30w WD controls. **C**, shows after tamoxifen at 6 to 8 weeks of age mice were placed on a Western diet (WD) for 18 weeks and then switching mice to chow diet for 12 weeks and then with IgG for 8 weeks on CD (18w WD+12w CD+8w IgG on CD). **D**, Shows after tamoxifen at 6 to 8 weeks of age mice were placed on a Western diet (WD) for 18 weeks and then switching mice to chow diet for 12 weeks and then treated with IL-1β Ab for 8 weeks on CD (18w WD+12w CD+8w IL-1β on CD) increased lesion size, increased necrotic core area, and increased cholesterol crystals compared to age and time of diet matched IgG controls. Graphical Images were created with BioRender.com

## Discussion

Atherosclerosis is a chronic inflammatory disease^30^ whose clinical complications, including myocardial infarction (MI) and stroke, are the leading causes of death worldwide^31^. Given the compelling evidence that inflammation plays a key role in development of atherosclerosis^7^, the expectation was that potent suppression of inflammation combined with aggressive lipid lowering would markedly reduce late-stage disease complications. Indeed, the CANTOS clinical trial testing an IL-1β Ab, Canakinumab, provided compelling evidence validating the inflammation hypothesis of atherosclerosis^12^. Moreover, in certain subsets of patients Canakinumab treatment reduced CV deaths by a remarkable 31%^13^. However, it failed to get FDA approval and the study leaves uncertainties regarding mechanisms influencing the response to IL1β antibody treatment and how to identify individual patients who would achieve sufficient benefit relative to the small increase in the risk of lethal infections. There is extensive ongoing work in this area mainly focused on the defining the role of clonal hematopoiesis and increased IL1β expression by macrophages as well as identifying genetic determinants that influence the response of lesion cells to IL1β^32,33^. The results of the present studies and our 2018 *Nature Medicine* paper^17^ may provide further insight in this area in that we unexpectedly found that IL-1β has both beneficial and detrimental effects in the pathogenesis of advanced atherosclerotic lesions. Indeed, we found that treatment of SMC lineage tracing *Apoe^−/−^*mice that had advanced lesions with a murine IL-1β Ab had multiple detrimental effects including a >50% reduction in SMC number and collagen content within the fibrous cap. However, we also showed that SMC-specific knockout of *IL1r1* was associated with both beneficial and detrimental effects including a >70% decrease in lesion size but a marked reduction in SMC investment into the protective fibrous cap.

Results of the present studies extend our previous studies by showing that IL-1β also has an essential role in mediating increases in indices of plaque stability in response to diet-induced reductions in lipids. Indeed, to our complete surprise, we found that IL-1β inhibition resulted in multiple detrimental changes including: 1) a 50% increase in lesion size; 2) significantly higher cholesterol crystal accumulation, intra-plaque hemorrhage, and increased necrotic core area, plaque burden and senescence; 3) reduced ACTA2^+^ cap thickness; and 4) increased SMC-derived lesion cells exhibiting an osteochondrocyte phenotype. Moreover, we recently showed that endothelial specific knockout of *IL1r1* unexpectantly resulted in an increase not a decrease in lesion size as well as a reduction in EC contribution to the ACTA2^+^ fibrous cap via EndoMT^3^. Our results in mice are consistent with a number of previous studies showing that pro-inflammatory signaling may have beneficial or detrimental effects on the pathogenesis of atherosclerosis in different cell types or even within a given cell type at different stages of disease progression^34,35^. As such, there is a need to identify more nuanced approaches for inhibiting the adverse effects of chronic inflammation without eliminating evolutionarily conserved beneficial functions essential for tissue repair, immune resistance to pathogens, and inflammation resolution.

The possible mechanisms by which IL-1β Ab treatment of mice after chow diet-induced lipid lowering resulted in extensive cholesterol crystal accumulation and failed reductions in lesion size are unclear. However, results of our present study suggest that it involves loss of IL-1β-mediated responses in multiple cell types as follows. Cholesterol crystal formation serves as both a priming and danger signal to activate the inflammasome resulting in activation of caspase1 and generation of IL-1β by macrophages^36^. Although macrophages are the primary source of IL-1β, other lesion cell types including SMC, EC, neutrophils, and T cells are the main lesion cells that respond to it. The effects of IL-1β on these cell types include it inducing: a pro-inflammatory state in SMC^37^; activation of leucocyte adhesion receptors in EC^38^; NET formation by neutrophils^39,40^; and T cell upregulation of IL-17^41,42^ which drives the chemokines CXCL1 and CXCL2 to promote further neutrophil recruitment^36^ (**summarized in a graphical abstract supplemental Figure XIV**). We postulate that these evolutionarily conserved IL-1β dependent protective mechanisms act cooperatively to remove cholesterol/cholesterol crystals and apoptotic cells but are over-whelmed and lesions continue to progress as long as mice continue on a WD. Upon switching to a low-fat chow diet these IL-1β-dependent processes are now sufficient to result in reduced lesion size and cholesterol content as well as increased plaque stability (**Graphical abstract in supplemental Figure XIV**).

The results of our studies herein provide clear evidence that inhibiting IL-1β negates many of the beneficial effects of dietary reductions in lipids in *Apoe^-/-^* mice with advanced atherosclerotic lesions. However, in contrast to these results, the results of the CANTOS clinical trial indicate that canakinumab (IL-1β Ab) treatment may be an appropriate therapy for subsets of high-risk post-myocardial infarction subjects^13^. It is easy to reconcile these apparent discrepant results as another case of mouse studies being a poor model of human disease due to inherent differences between humans and mice, disease time-course, treatment periods, immune system differences and experimental differences including effects of prior myocardial infarction, and drug treatments not present in the mouse model. However, two studies (MRC-ILA Heart Study^43^ and IL-1 Genetics Consortium^44^) suggest that inhibition of IL-1 signaling may unexpectedly increase the risk for some cardiovascular events. Moving forward, it will be critical to determine strategies to identify subsets of patients that will exhibit the most benefit from specific anti-inflammatory therapies and to determine whether there are patients more susceptible to adverse effects. Consistent with this possibility, Fuster et al. found that mutations in hematopoietic stem cells could affect atherosclerosis development by modifying the function of myeloid cells^32^. One prevalent mutation of Tet2 exacerbated atherosclerosis in LDL receptor KO mice and was accompanied by enhanced IL-1β production by macrophages^32^. Personalized approaches to cardiovascular care targeting pro-atherogenic clonal hematopoiesis with anti-inflammatory medications may be possible. The greatest value of the mouse studies is to generate hypotheses for further testing in humans to better understand the mechanistic and physiological aspects of inhibition of inflammation and to then identify alternative or combinatorial anti-inflammatory therapeutic targets that are both effective and safe. Although animal models can be valuable for testing hypotheses derived from human studies and clinical trials, there is the need to better match animal models to the cardiovascular disease and clinical need of interest, and for investigators to perform more human validation studies.

Given the prevalence of lipid-lowering medication in the treatment of cardiovascular disease, we believe the use of lipid-reduction mouse models are critical to the testing and understanding of future intervention therapies. The CANTOS trial was successful in demonstrating that reducing inflammation markedly improves cardiovascular outcomes in select individuals. As such, the rest of the story will rely on having an in-depth understanding of the impact of these therapies on the cellular components of atherosclerotic lesions. Indeed, targeting inflammation is undoubtedly a key component of eliminating the scourge of cardiovascular disease from humanity. However, we believe that accomplishing this goal will require identification of therapeutic approaches customized for the individual patient and which selectively target detrimental and not beneficial inflammatory responses.

## Acknowledgments

We thank the University of Virginia Advanced Microscopy core, Flow Cytometry Core, and Genome Analysis and Technology Core (RRID: SCR_018883). We also thank Novogene for assistance with the bulk-RNA sequencing experiments. Thanks to Dr. Gerry Waters, liaison to Novartis Institutes for Biomedical Research, for providing the IL-1β Ab and IgG control antibodies as well as valuable recommendations regarding the manuscript presentation..

## Author Contributions

SK conceptualized, designed, and performed the bulk of the experiments, validation, data collection, analysis and interpretation, and contributed significantly to development of methodology, manuscript writing and editing. VK contributed to data collection and interpretation, analysis of indices of stability-movat, picrosirius analysis, IFC staining, confocal imaging, counting the single cells of confocal images, and sample preparation for mouse bulk RNA-seq and help in harvesting mice and collecting samples and data interpretation and manuscript editing. RAD contributed to data interpretation and manuscript editing. LSS contributed to data analysis and data interpretation and manuscript editing. CMW contributed to staining the antibodies for Cy-TOF data collection, and data analysis. EB contributed to data collection and analysis. XB contributed to data collection and analysis. GFA contributed to analysis of scRNA-seq data (bioinformatics part). GBB, and SK contributed to data collection and analysis. RAB contributed to data interpretation and manuscript editing. ERZ contributed to Cy-TOF data collection and analysis. HMR and GP contributed to data analysis and interpretation. GKO supervised the entire project, conceptualization, data interpretation, funding and manuscript writing and editing.

## Supplemental Methods

### Plaque burden analysis by Sudan IV staining

Aortas from *Myh11-CreERT2 R26R-eYFP Apoe^−/−^* progression, regression and mice treated with anti-IL-1β antibody or IgG2a control for 8 weeks were dissected from the aortic arch to the iliac bifurcation and subjected to en face Sudan IV staining^17^. In brief, after carefully removing the peri-adventitial fat, aortas were dehydrated in 70% ethanol for 5 minutes and incubated in Sudan IV solution for 6 minutes, which was prepared as follows: 1g of Sudan IV (Sigma Aldrich, S4261) was diluted in 100 mL of 70% ethanol and 100 mL of 100% acetone. Finally, aortas were differentiated in 80% ethanol for 3 minutes and stored in PBS at 4°C, and photographs were taken with a mobile phone camera.

### Senescence-Associated β–Galactosidase (SAβG) Staining

SAβG staining for senescence on aortas were examined using the senescence detection assay kit (Millipore-Sigma Cat No.KAA002, Rockville, MD 20850, USA)^45^. Freshly isolated aortas were kept on ice in a 12 well plate containing PBS until all the aortas were harvested. Aortas were washed twice with the PBS, and then fixed with the 1x diluted fixation solution provided in the kit for 10 minutes as per the manufacturer’s instructions. After the fixation, aortas were washed thrice with PBS and then 1mL of freshly prepared SAβG solution was added to a well of 12 well plate containing one aorta per well. Then, plates were wrapped with aluminum foil to avoid light exposure and placed in a 37^0^C incubator for 24 hours. After that, aortas were washed thrice with PBS, and photographs were taken with a mobile phone camera.

### Immunohistochemistry and atherosclerotic plaque morphometry

Paraformaldehyde-fixed paraffin-embedded BCAs were serially cut into 10 μm thick sections from the aortic arch to the bifurcation of the right subclavian artery. For morphometric analysis, we performed modified Russell-Movat staining on BCA as previously described^3,17^. Picrosirius Red staining were performed to assess collagen content on BCAs^3,17^. Immunohistochemistry (IHC) was performed with antibodies against Ter-119 (SC-19592-1μg/mL, Santa-Cruz)^3,17^. Staining for immunohistochemistry was visualized by DAB (Acros Organics).

### Immunofluorescent staining

BCA sections were de-paraffinized and rehydrated in xylene and ethanol series. After antigen retrieval (H-3300-250, Vector Laboratories, Newark, CA 94560, USA), sections were blocked with fish skin gelatin–PBS (6 g/L) containing 10% horse serum for 1 hour at room temperature. Slides were incubated with the following antibodies: goat polyclonal anti-GFP^3,17,23^ (4 μg/mL, ab6673, Abcam, Waltham, MA 02453, USA) for detection of eYFP (Overnight 4^0^C) and Lgals3^3,17,23^ (CEDARLANE-CL8942AP) (Overnight 4^0^C). Mouse monoclonal SM α-actin-Cy3 (Acta2)^3,17,23^ (4.4 μg/mL, clone 1A4, C6198, Sigma Aldrich, Rockville, MD 20850, USA) 1hr at room temperature. Secondary Abs Donkey anti-goat conjugated to Alexa 488^3,17,23^ (A11055-5 μg/mL, Invitrogen, Waltham, MA 02451, USA.), and donkey anti-rat conjugated to Dylight 650^24^ (SA5-10029 5μg/mL, Invitrogen, Waltham, MA 02451, USA.). DAPI^23^ (0.05 mg/mL, D3571, Thermo Fisher Scientific, Waltham, MA 02451, USA.) was used as a nuclear counterstain, and slides were mounted using Prolong Gold Antifade^3^ (Invitrogen.P36930, Waltham, MA 02451, USA.).

### Imaging

Movat, Ter119 staining of brachiocephalic arteries were imaged using a thunder imager Leica microscope. Image acquisition was performed with Leica software. The lumen, lesion, necrotic core areas, cholesterol crystals as well as the external elastic lamina (vessel area), were measured on digitized images of the Movat staining and collagen content (picrosirius red) using Fiji software version 1.53c. Immunofluorescent staining was imaged using a Zeiss LSM880 confocal microscope to acquire a series of z-stack images at 1-μm intervals. Zen 2009 Light Edition Software (Zeiss) was used for analysis of each z-stack image, and single-cell counting was performed for phenotyping and quantifying the cell population comprised within the 30μm thick layer proximal to the lumen (i.e., fibrous cap area)^3^. Assessment of ACTA2+ cap thickness (normalized to lesion) was performed using Zen 2009 Light Edition Software^3^. Maximal intensity projection of representative images was used to generate the representative images included in the figures.

### ScRNA-seq on micro dissected BCA lesions and analysis

We used a similar protocol as we described in a recent article^24^.

***Micro-dissection of BCA lesions and Cell processing for scRNA-seq:*** Dissection Opened mouse and gravity perfused with PBS+ 1ug/mL actinomycin D (Gibco, #11805017). Cleaned BCA and slice open from carotid/subclavian bifurcation toward aortic root. Peeled off advanced BCA lesion (without media) using fine forceps. BCA plaques were placed in Lo-bind tube with 25 µL FACS buffer (1% BSA in PBS) on ice. ***Digestion:*** 500 µL of digestion buffer (50mg/10mL Liberase (Roche, #355374) + 3.5 ug/mL elastase Worthington # 2279 35H8143 in RPMI +1 ug/mL actinomycin D) was added and lesion samples were chopped with scissors. Samples were incubated in an incubator for 1 hr at 37^0^C. ***Centrifugation/Filtration:*** After digestion, 1 mL of FACS buffer (1% BSA) was added to samples and filtered through 70 um strainers into an eppendorf tube coated with FACS buffer. Centrifuged at 1000xg for 10 min at 4^0^C. Removed supernatant and using Lo-bind pipet tip, cell pellet was suspended in 250 µL 0.04% non-Acetylated BSA (ThermoFisher #AM2616). ***Sorting:*** Samples were stained with sytox blue (Life technologies #S11348) viability dye and sorted with a BD FACS influx cell sorter with gating on live, single cells that were eYFP+. Cells were collected into Lo-bind tubes containing 0.04% non-acetylated BSA. ***Sequencing:*** Chromium 10X GEM bead prepared by UVA Genome analysis and technology core facility. Briefly, two thousand cells in each group were targeted in Chromium 10X genomics libraries, which, after barcoding, were pooled and sequenced on the Illumina, 150 cycle high-output. Gene-barcode matrices were analyzed in R using Seurat v3. Cells were filtered for 200 to 5,000 reads per unique molecular identifiers, ≤10% mitochondrial and <5% hemoglobin gene content. Differential expression analysis was done using Model-Based Analysis of Single-Cell Transcriptomics (MAST). Code is available on request. scRNA-seq performed at Next Seq Genome Analysis and Technology Core RRID: SCR_018883, University of Virginia.

### Bulk RNA-seq on micro dissected BCA lesions and Data Analysis

Total RNA was isolated using RNeasy kit (Qiazen 74106) from the micro-dissected BCA lesions without media of *Myh11-CreERT2 R26R-eYFP Apoe^−/−^* mice fed a high-fat Western diet for 18 weeks plus 20 weeks of chow diet and treated with the anti-IL1β antibody or with the IgG control antibody for the last 8 weeks of standard chow diet feeding (that is 31 to 38 weeks). Total RNA was also isolated from the micro-dissected BCA lesions of progression and regression *Apoe^−/−^* mice. The RNA library was prepared according to manufacturer instructions with rRNA reduction with an average RNA integrity number (RIN) of 6.6. RNA libraries were provided to the company Novogene Inc. (USA) for bulk RNA-seq. Libraries were sequenced with the Illumina short-read sequencing (IlluminaHiSeq v4; 100 bp and 25 million paired-end reads). Raw reads were QCed to exclude reads containing adapters, reads containing ambiguous calls (N) >10%, and reads where 50% of bases had a Qscore <5. QCed reads were mapped to the Mus musculus mice reference genome using the STAR software. Reads per transcript were extracted from the STAR output and converted to FPKM (Fragments per Kilobase of transcript sequence per Million mapped reads). 54,532 transcripts were obtained, of which 21,116 transcripts had an average expression >1 read per sample and were used for differential gene expression. Read counts were normalized and differential gene expression between groups was performed using DESeq2 in R53. Reactome gene enrichment analysis for genes with significant up-or down-regulation was performed.

### CyTOF sample preparation

Cell suspensions were prepared as described for scRNA-seq. Samples were washed with PBS, treated with 5μM cisplatin (Sigma Aldrich) for 30 seconds, quenched with Cell Staining Medium (CSM) (0.5% w/v Fraction V BSA (VWR) in PBS with 0.02% w/v sodium azide (Sigma Aldrich), washed again with PBS (Gibco), and fixed in 1.6% paraformaldehyde (Electron Microscopy Sciences) for 10 minutes with intermittent vortexing. Cells were washed into CSM and stored at −80^0^C. Samples were barcoded with palladium reagents and pooled. The samples were washed with CSM, and a single surface antibody cocktail (supplemental table) was prepared and added to the barcoded samples incubating at room temperature with shaking for 30 minutes. Barcoded samples were again washed with CSM and permeabilized with −20C 100% methanol (VWR), incubating for 10 minutes on ice and vortexing intermittently. After washing with CSM, samples were incubated in an intracellular antibody cocktail^23^ at room temperature with shaking for one hour. Samples were washed in CSM and incubated overnight at 4C with 1:5000 Iridium intercalator (Fluidigm) in 1.6% paraformaldehyde in CSM. Samples were then washed with CSM and washed into water with normalization beads^46^ and run on a Helios 2 mass cytometer (Fluidlike).

### CyTOF data analysis

After measurement the samples were normalized^46^, de-barcoded^47^, and uploaded to Cytobank (www.cytobank.org) for cleanup gating before exporting FCS files for downstream analysis. Raw data was transformed using a division factor of 5^48^. Dimensionality reduction was done by UMAP using default parameters before using the full graph output by UMAP for Leiden clustering with default parameters.

