## Supplementary material for "IL-1β inhibition partially negates the beneficial effects of diet-induced lipid lowering": supllemental figures

Short Title: IL-1β inhibition reduces benefits of lipid lowering.

[1]Robert M. Berne Cardiovascular Research Center, University of Virginia, Charlottesville, USA.

[2]Laboratory of Experimental Cardiology, University Medical Center Utrecht, Utrecht University, the Netherlands.

*** Corresponding author:**

Dr. Gary K. Owens

Univ. of Virginia School of Medicine

Robert M. Berne Cardiovascular Research Center

PO Box 801394

MR5 Building

Charlottesville, Virginia 22908-1394

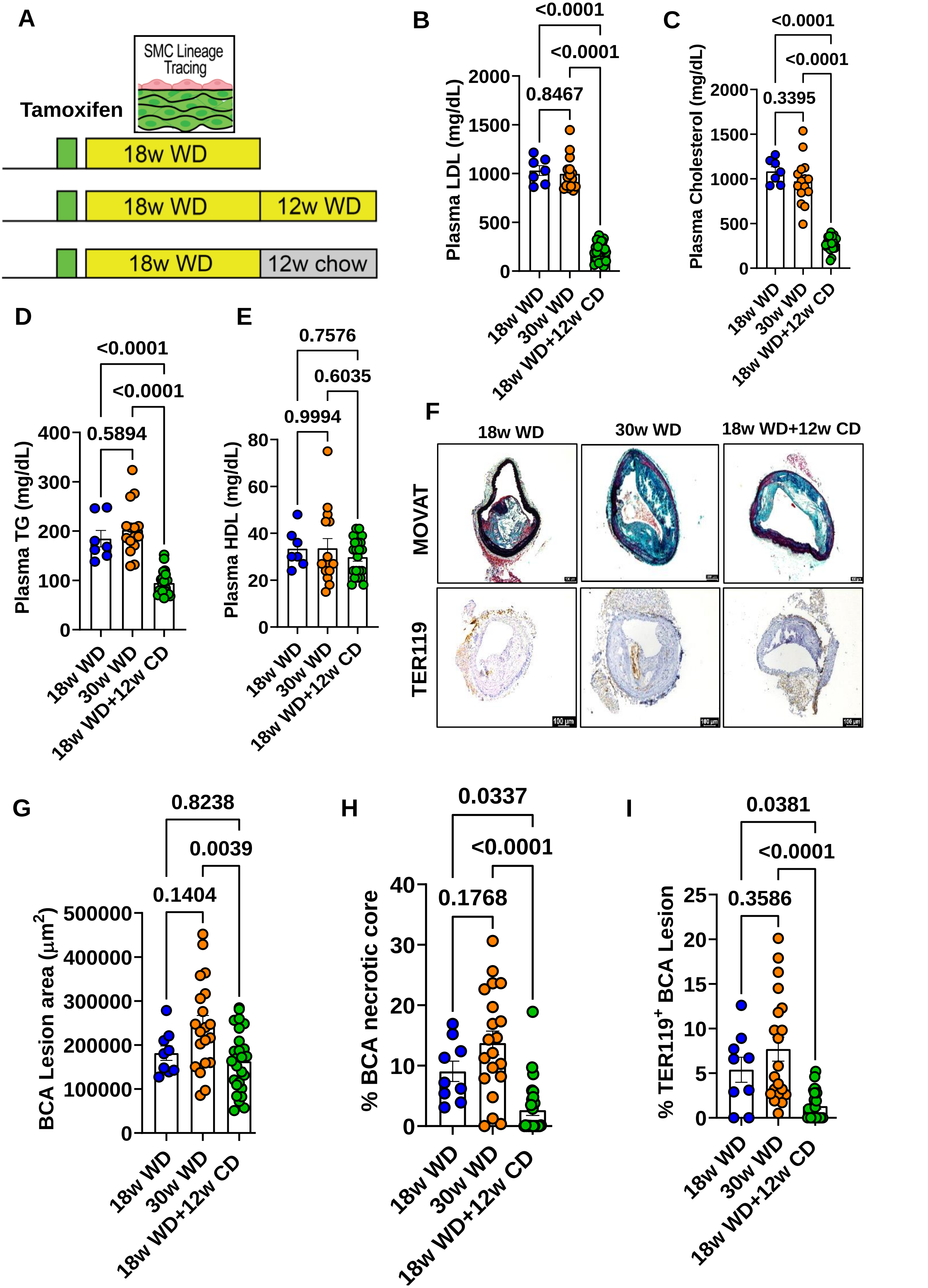


Supplemental Figure I

**Supplemental Figure I.** ***Apoe$^{-/-}$*** **mice fed a WD for 18 weeks followed by 12 weeks of a low fat chow diet show marked reductions in plasma LDL, lesion size, necrotic core area, and intraplaque hemorrhage as compared to age-matched mice that continue on a WD. A**, Experimental design, SMC lineage tracing *Apoe$^{-/-}$* mice were injected with tamoxifen at 6 to 8 weeks of age and subsequently placed on a Western diet (WD) for 18 weeks to induce advanced atherosclerosis and randomly distributed into two groups. One group continued receiving WD for another 12 weeks, and another group received chow diet for 12 weeks. **B**, plasma LDL concentration. **C**, plasma cholesterol concentration. **D**, plasma TG concentration. **E**, plasma HDL concentration. **F**, representative Movat, TER119 (intraplaque hemorrhage detection), stained BCA lesions. **G**, BCA lesion area. **H**, percentage BCA necrotic core area. **I**, percentage of TER119+ BCA lesions. Error bars represent mean ± SEM; p-values displayed refer to one-way ANOVA with multiple comparisons. Each circle on the graph indicates data from an individual mouse.

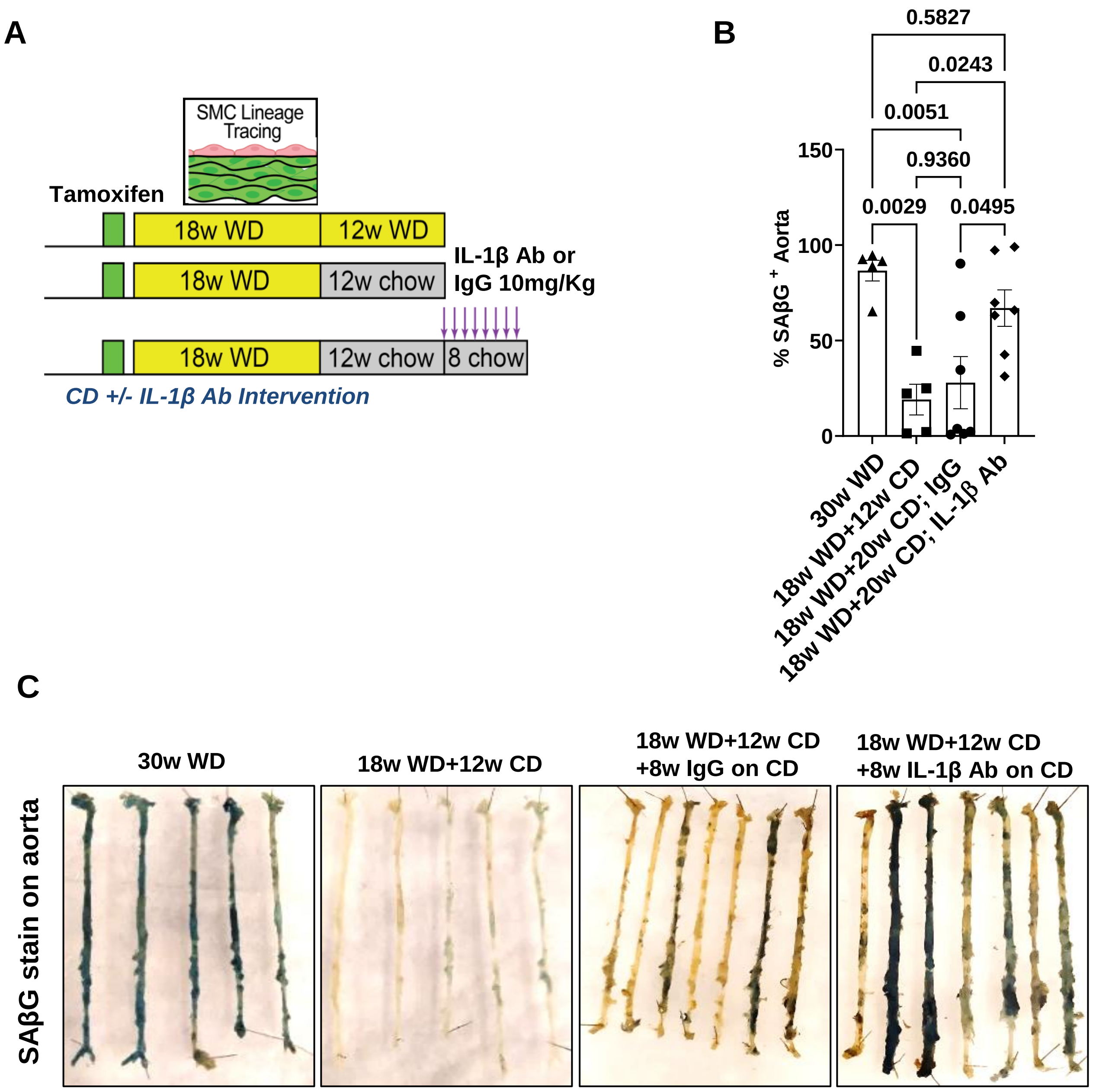


**Supplemental Figure II. Treatment of Apoe−/− mice with an IL-1β neutralizing Ab after chow diet-induced lipid lowering resulted in an increased senescence. A**, Experimental design, SMC lineage tracing Apoe−/− mice were injected with tamoxifen at 6 to 8 weeks of age and subsequently placed on a WD for 18 weeks to induce advanced lesion formation followed by switching mice to chow diet for 12 weeks or one group continued receiving WD for another 12 weeks as 30weeks WD progression group. Chow diet-switched mice were then randomized to being treated with a murine IL-1β neutralizing Ab or an isotype-matched IgG control Ab for 8 weeks while mice continued on chow diet. **B**, Quantification of SAβG+ area of aortas. **C**, SAβG (senescence marker) stained aortas. Error bars represent mean ± SEM, p-values displayed refer to one-way ANOVA with multiple comparisons. Each triangle, square, circle and diamond shape on the graph indicates data from an individual mouse.

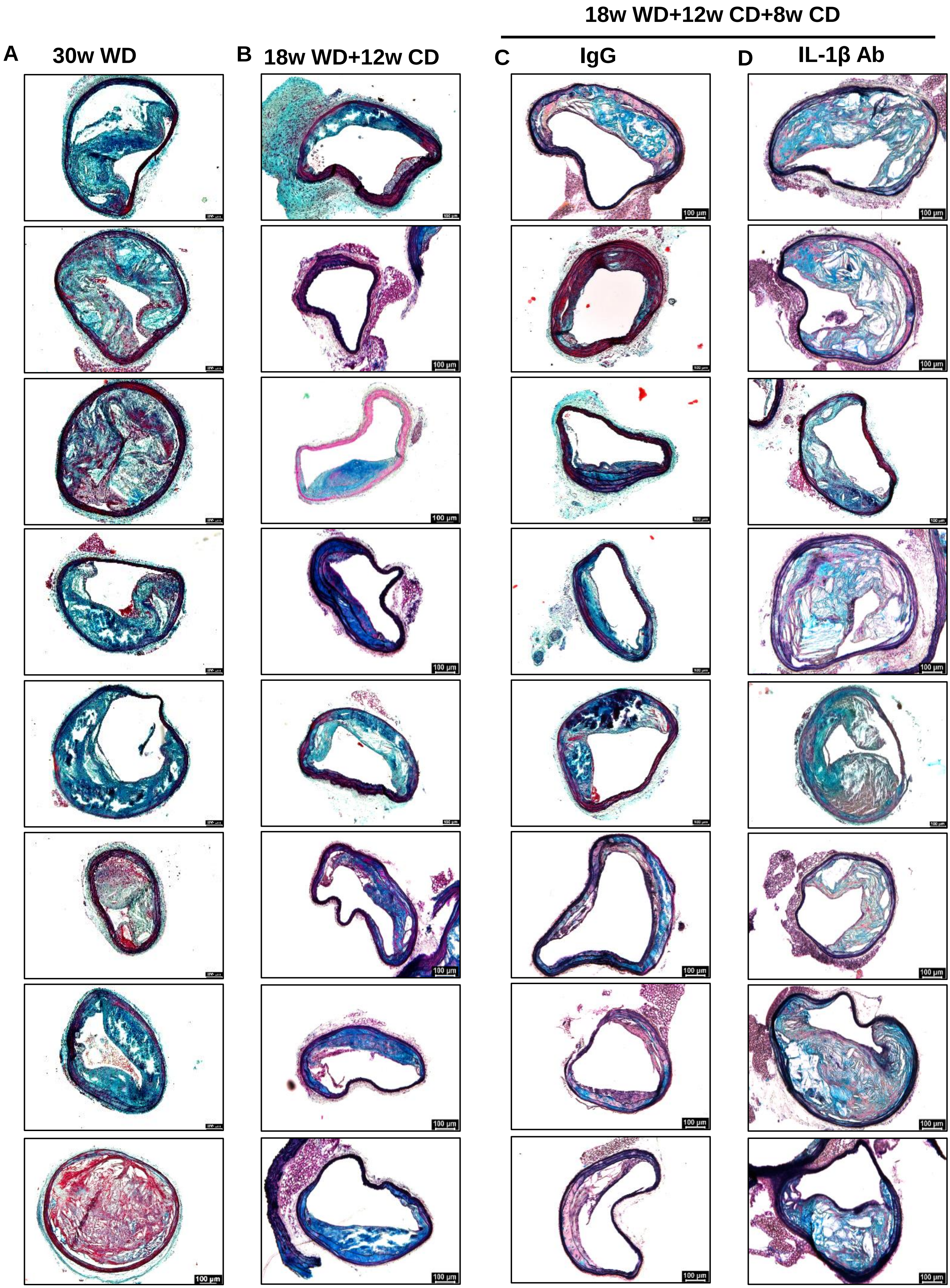


**Supplemental Figure III. Multiple representative BCA MOVAT images used in Figures 1F-1H analysis. A**, 30w WD. **B**, 18w WD+12w CD. **C**, 18w WD+12w CD+8w IgG on CD. **D**, 18w WD+12w CD+8w IL-1β Ab on CD.

### 30w WD; BCA lesions used in Figure 1F-1H

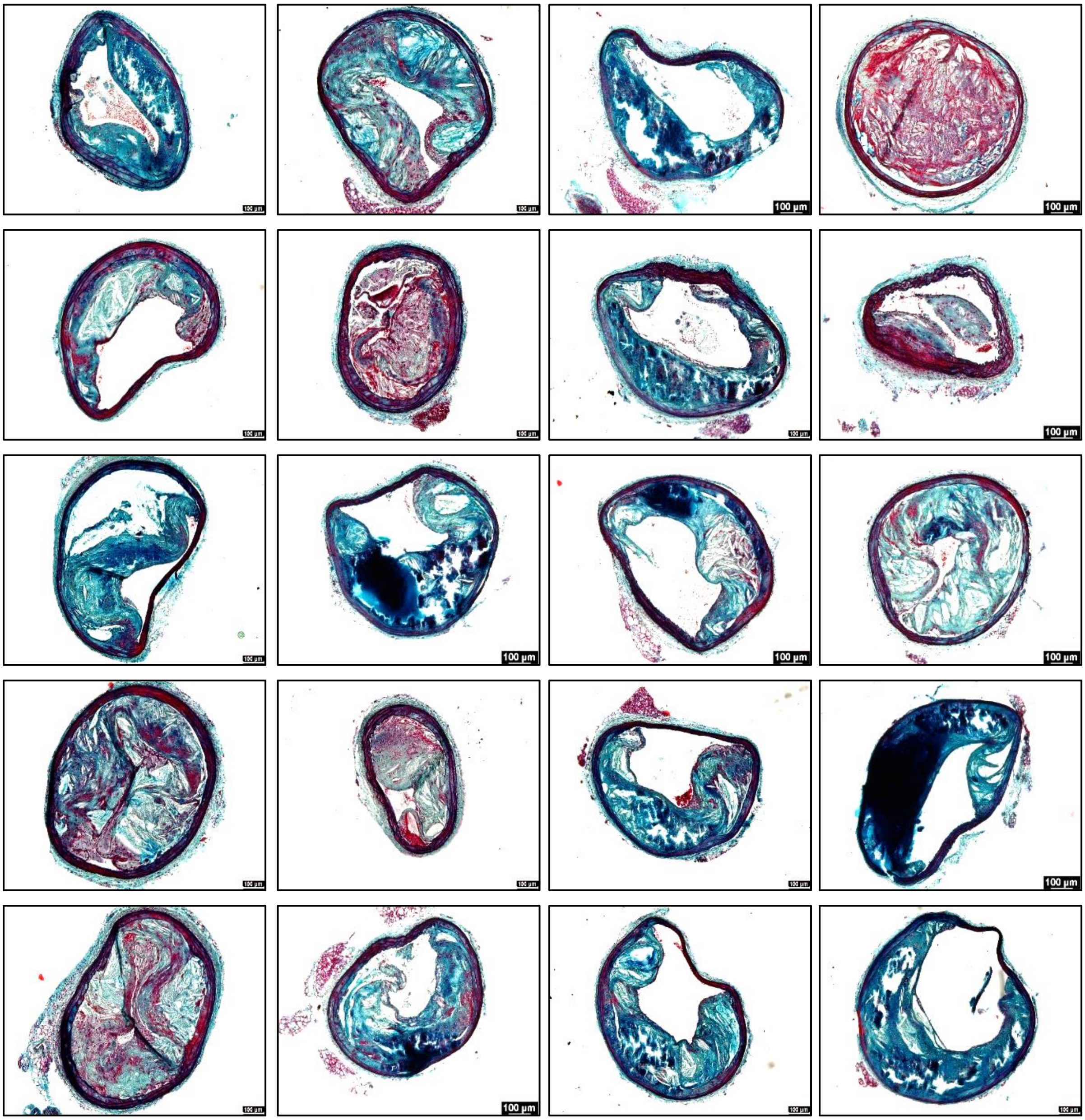

**Supplemental Figure IV. All the 30w WD fed mice BCA MOVAT images used in Figure 1F-1H analysis.**

### 18w WD+12w CD; BCA lesions used in Figure 1F-1H

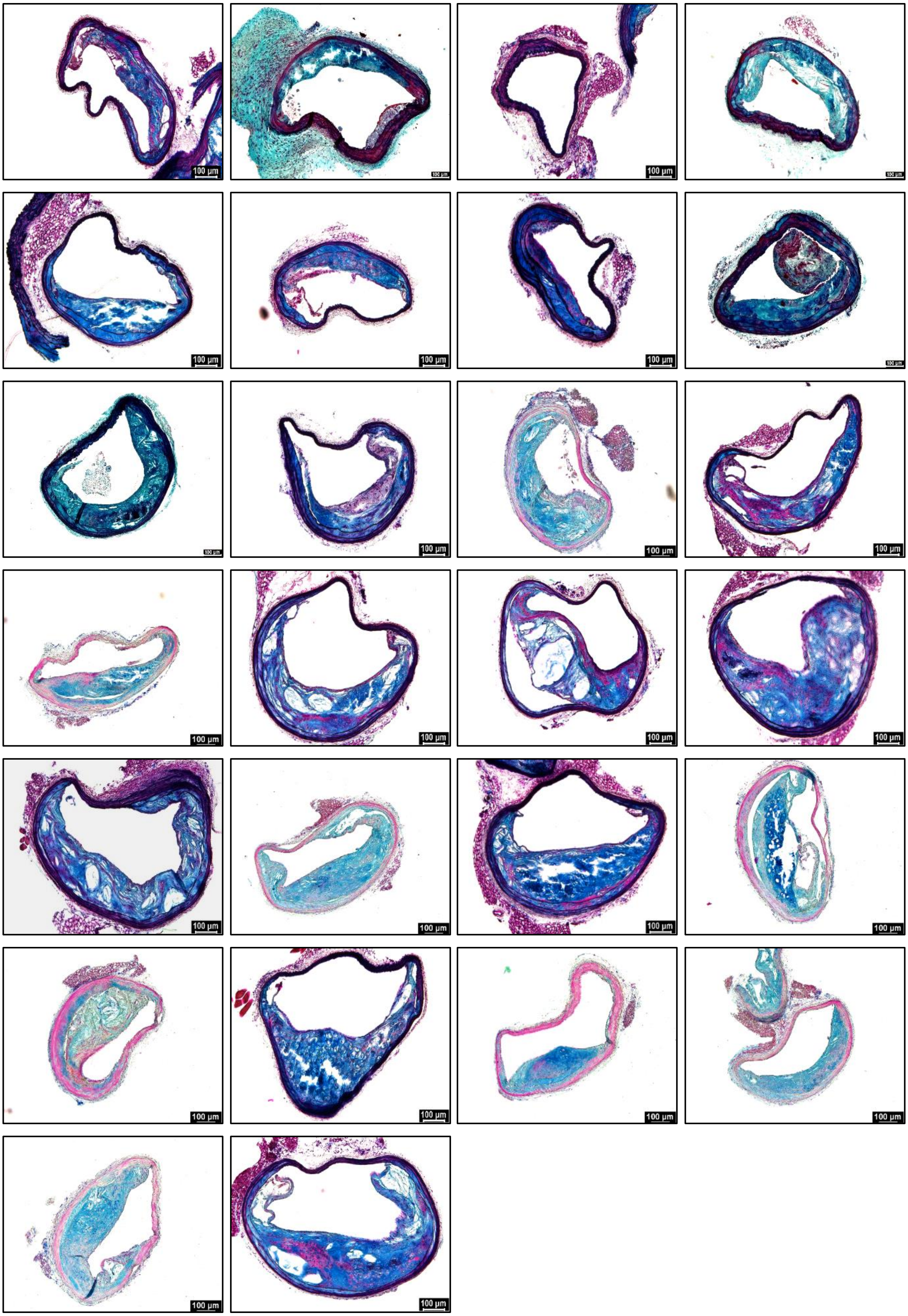

**Supplemental Figure V. All the 18w WD+12w CD fed mice BCA MOVAT images used in Figure 1F-1H analysis.**

### 18w WD+12w CD+ 8w IgG on CD; BCA lesions used in Figure 1F-1H

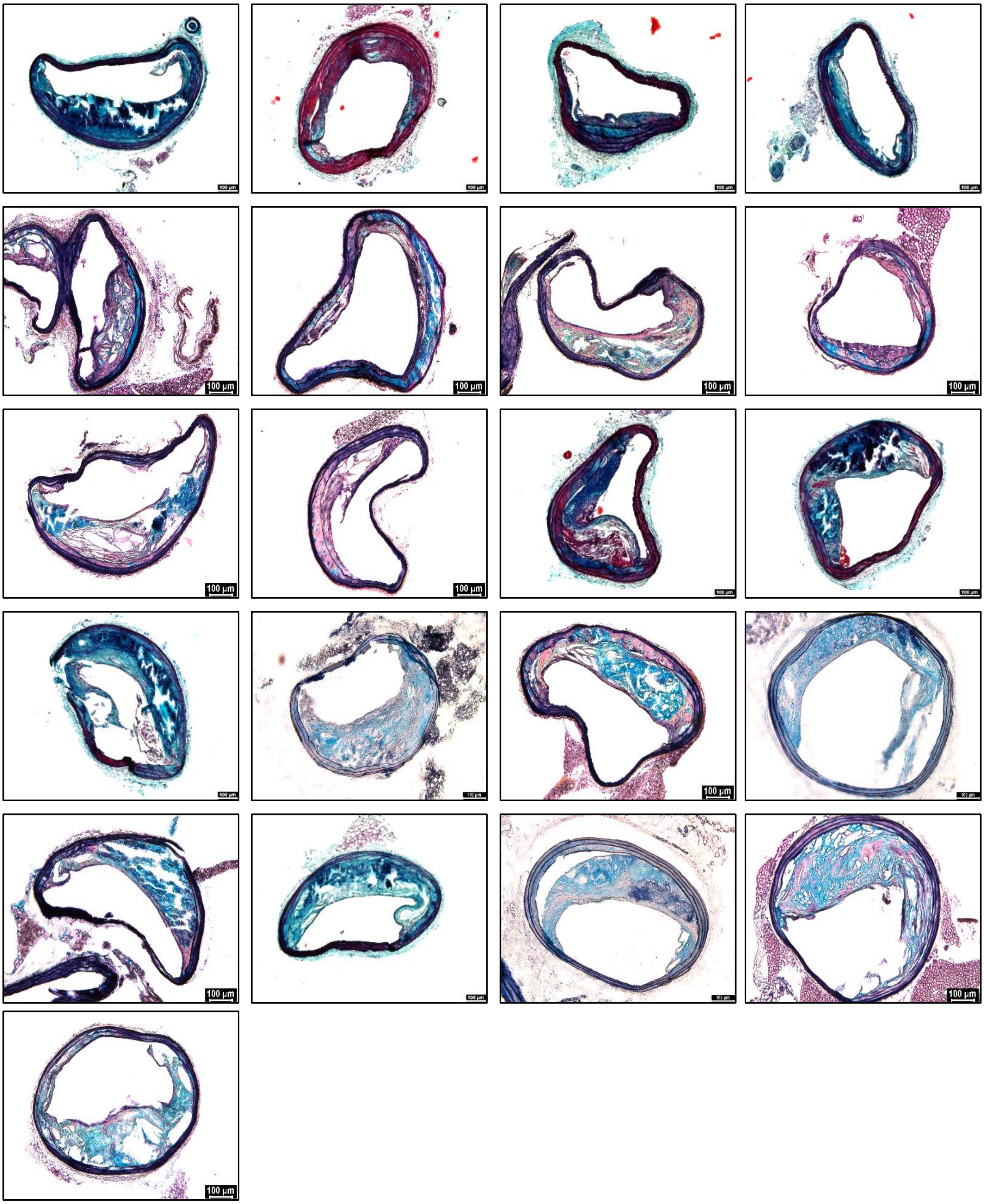

**Supplemental Figure VI. All the 18w WD+12w CD+8w IgG treated on CD group mice BCA MOVAT images used in Figure 1F-1H analysis.**

**Supplemental Figure VI**

**18w WD+12w CD+ 8w IL-1β on CD; BCA lesions used in Figure 1F-1H**

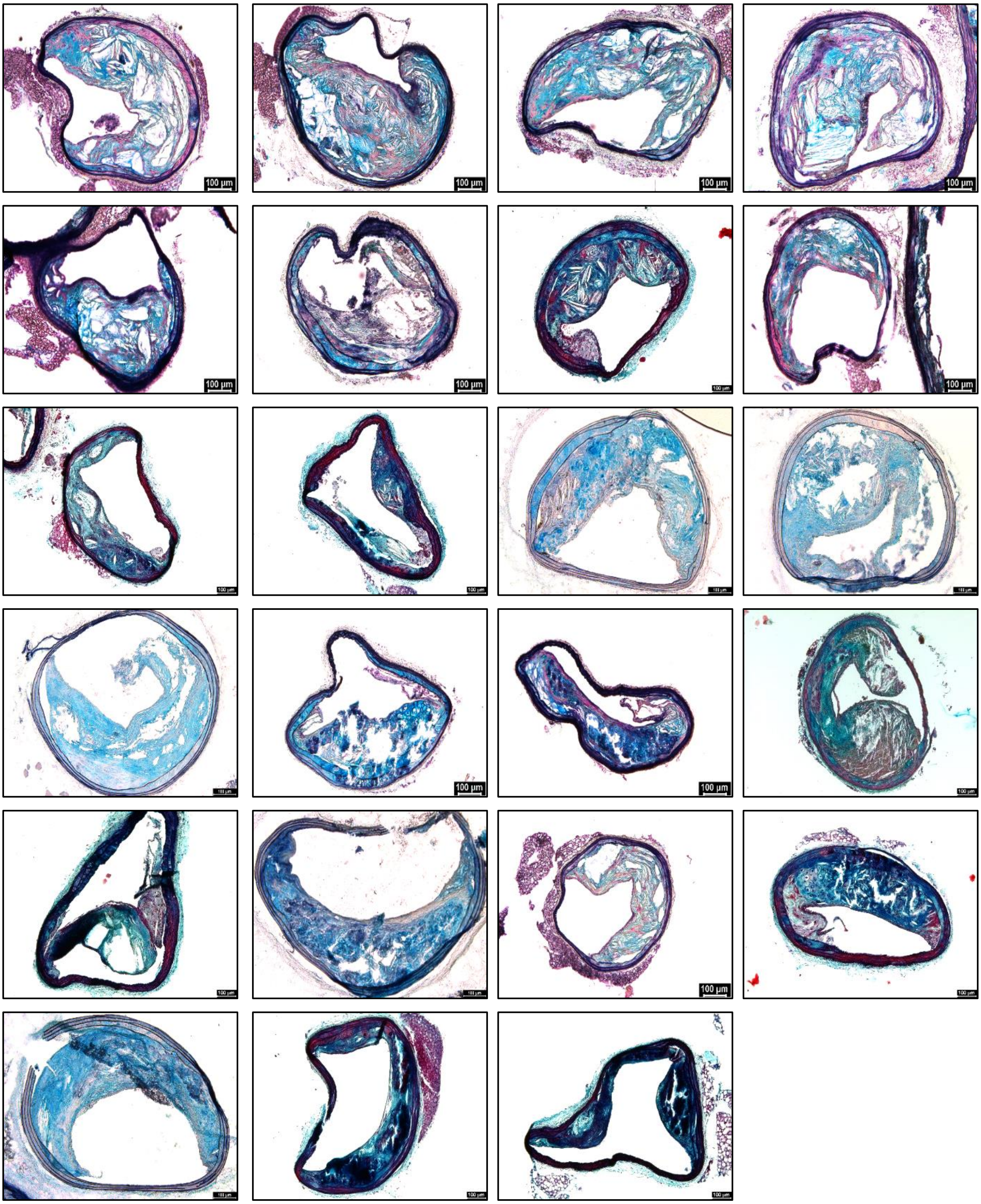


**Supplemental Figure VII. All the 18w WD+12w CD+8w IL-1β Ab treated on CD group mice BCA MOVAT images used in Figure 1F-1H analysis.**

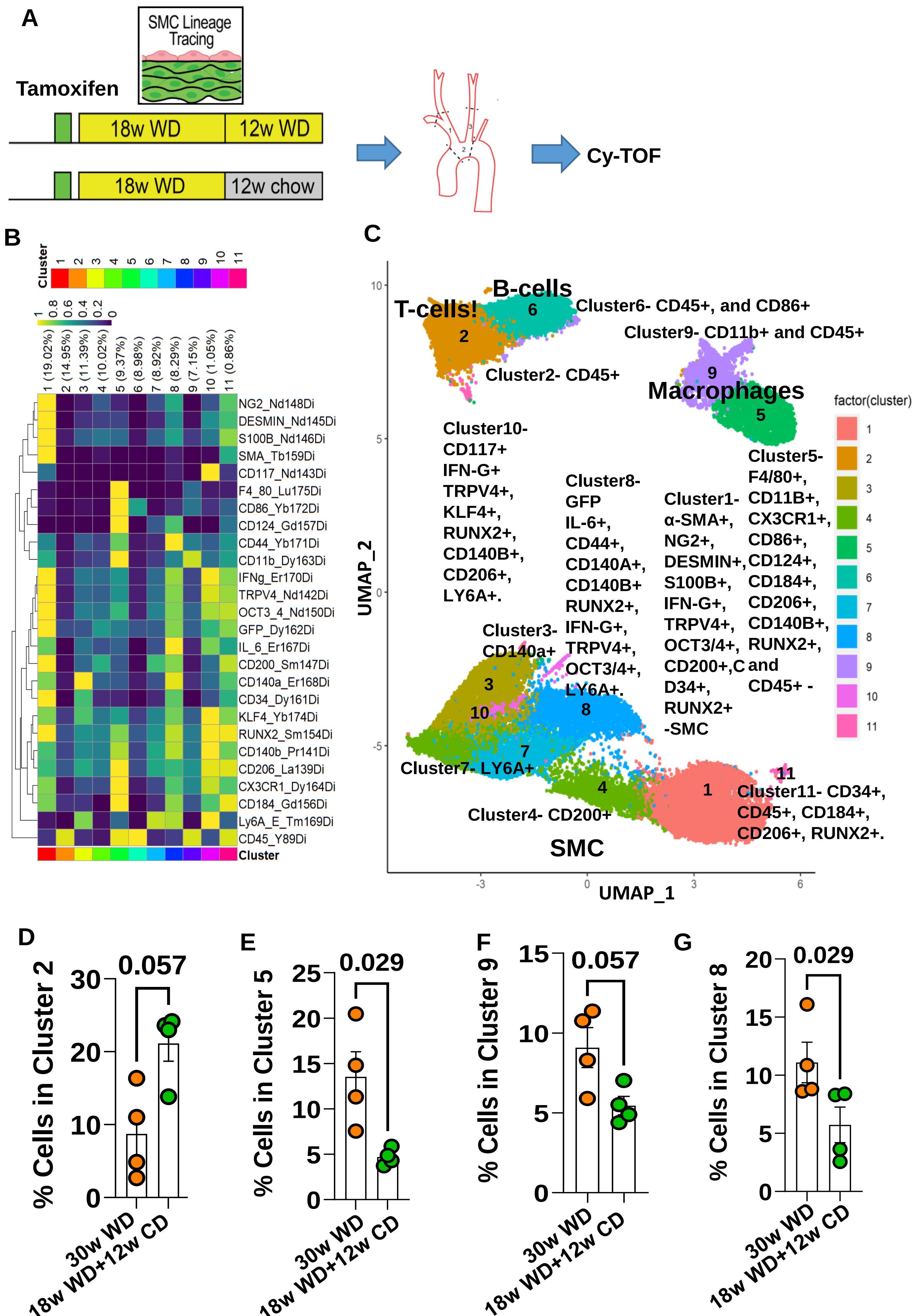


Supplemental Figure VIII

**Supplemental Figure VIII. Apoe−/− mice fed a WD for 18 weeks followed by 12 weeks of chow diet had reduced macrophages and osteochondrocyte cells in the aortic arch lesions including BCA and carotids compared to age-matched mice that continued on a WD. A**, Experimental design, SMC-lineage tracing *Apoe*$^{-/-}$ mice were injected with tamoxifen at 6 to 8 weeks of age and subsequently placed on a Western diet (WD) for 18 weeks to induce advanced atherosclerosis and randomly distributed into two groups. One group continued receiving WD for another 12 weeks, and another group received chow diet for 12 weeks. **B**, 26-antibody panel Cy-TOF staining on BCA region lesion cells including aortic arch area, BCA and carotids. **C**, UMAP colored by cluster from Cy-TOF analysis from B. **D**, percentage of cells in cluster 5 (macrophages). **E**, percentage of cells in cluster 9 (CD45+ and CD11b+ macrophage cells). **F**, percentage of cells in cluster 8 (osteochondrocyte/inflammatory markers+ cells). **G**, percentage of cells in cluster 2 (CD45+ but negative for the B-cell marker CD86). Error bars represent mean ± SEM, and p-values displayed refer to Mann-Whitney U-test between 30w WD and 18w WD+12w CD groups. Each circle on the graph indicates an individual mouse.

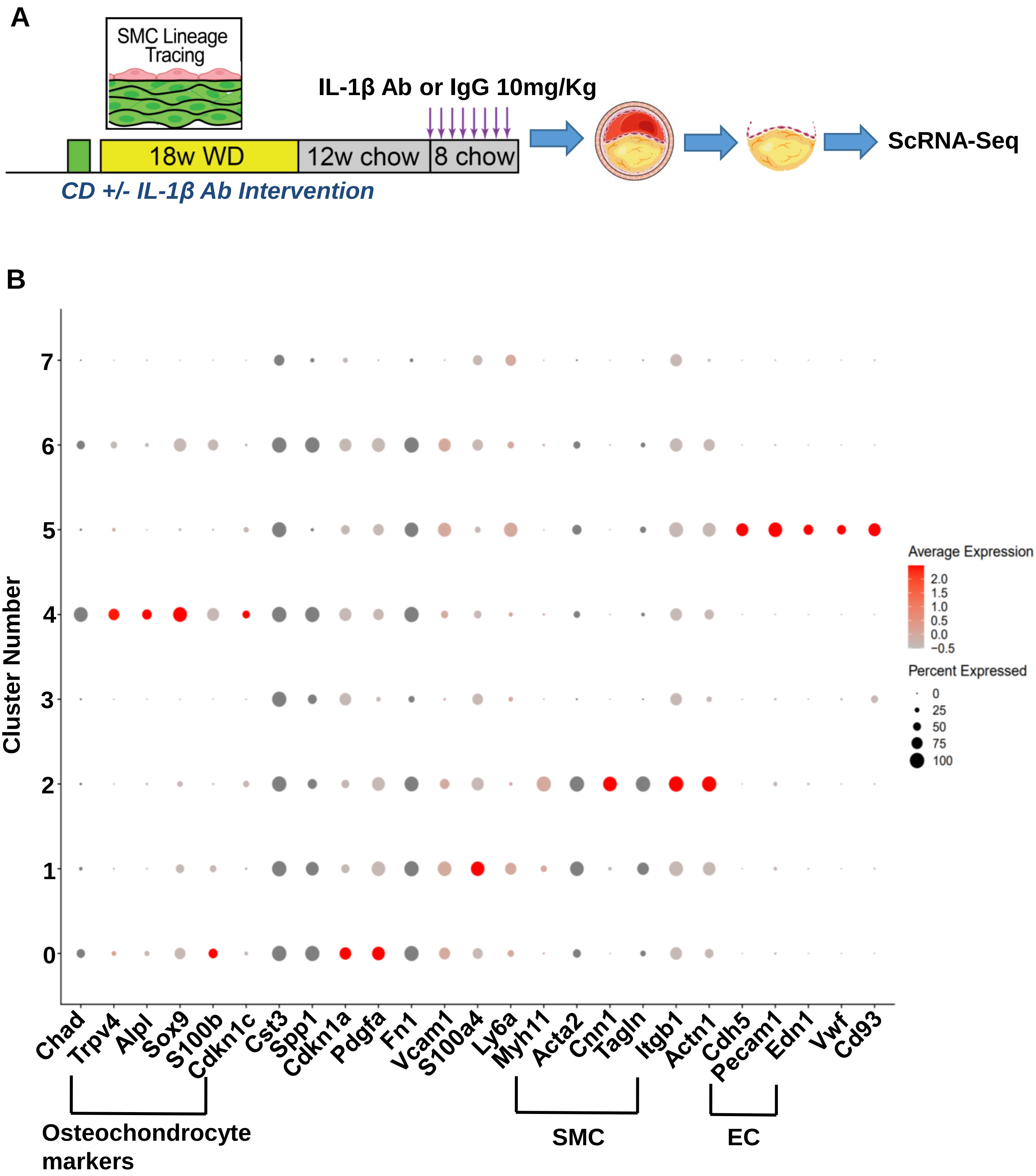


**Supplemental Figure IX. traditional markers to define the different cell types with a dot plot. A**, Experimental design, SMC-lineage tracing Apoe−/− mice were injected with tamoxifen at 6 to 8 weeks of age and subsequently placed on a Western diet (WD) for 18 weeks to induce advanced atherosclerosis and randomly distributed into two groups. One group continued receiving WD for another 12 weeks, and another group received chow diet for 12 weeks. Mice were then randomized to being treated with a murine IL-1β neutralizing Ab or an isotype-matched IgG control Ab for 8 weeks while mice continued on chow diet. **B**, markers to define the different cell types with dot plot.

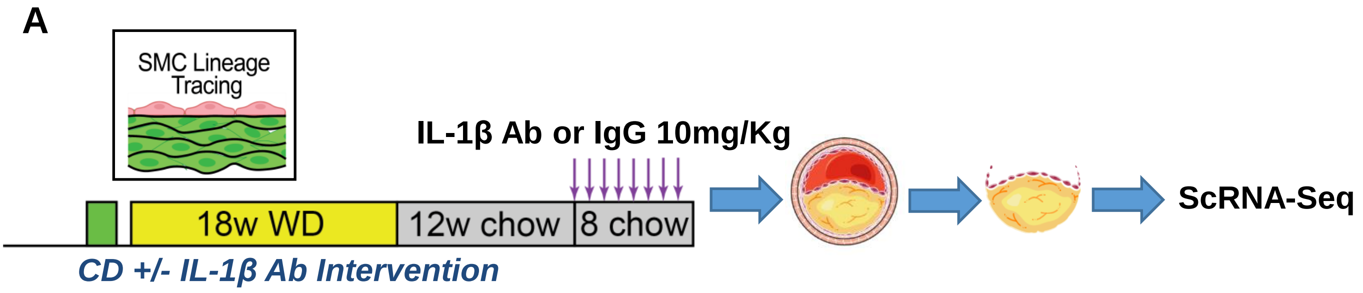


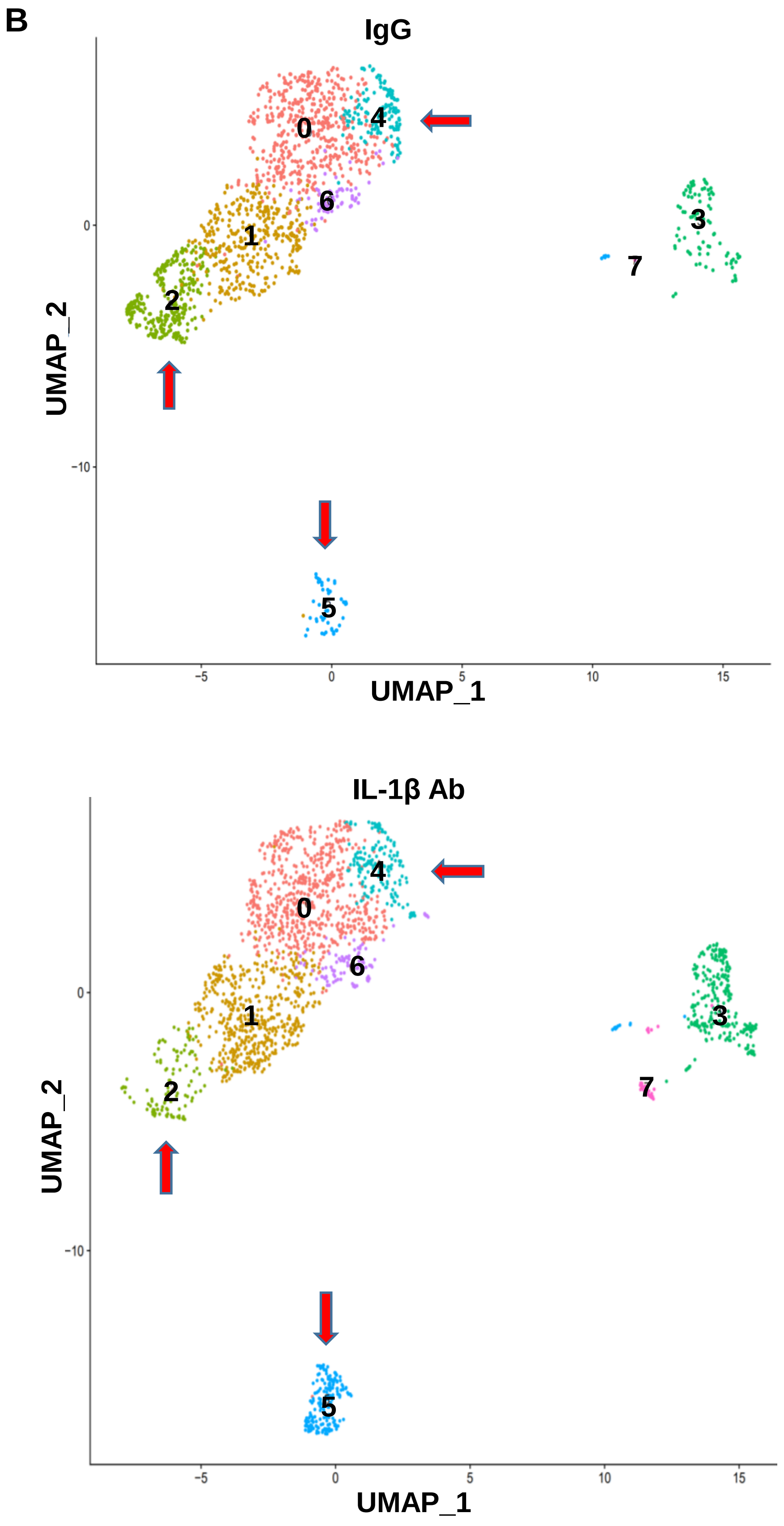


**Supplemental Figure X. IL-1β Ab Intervention in *Apoe*$^{-/-}$ mice after chow diet-induced lipid lowering resulted in a reduction in SMC marker+ clusters and an increase in the osteochondrocyte marker+ clusters of BCA lesions compared to age and diet matched IgG controls. A**, Experimental design, SMC-lineage tracing Apoe−/− mice were injected with tamoxifen at 6 to 8 weeks of age and subsequently placed on a Western diet (WD) for 18 weeks to induce advanced atherosclerosis and randomly distributed into two groups. One group continued receiving WD for another 12 weeks, and another group received chow diet for 12 weeks. Mice were then randomized to being treated with a murine IL-1β neutralizing Ab or an isotype-matched IgG control Ab for 8 weeks while mice continued on chow diet. **B**, UMAP analysis shows cells from IgG and from IL-1β Ab. Red arrows indicating number of cells were changed in cluster-2 (SMC marker+), cluster-4 (osteochondrocytes marker+) and cluster-5 (EC) of IL-1β Ab group compared to IgG group.

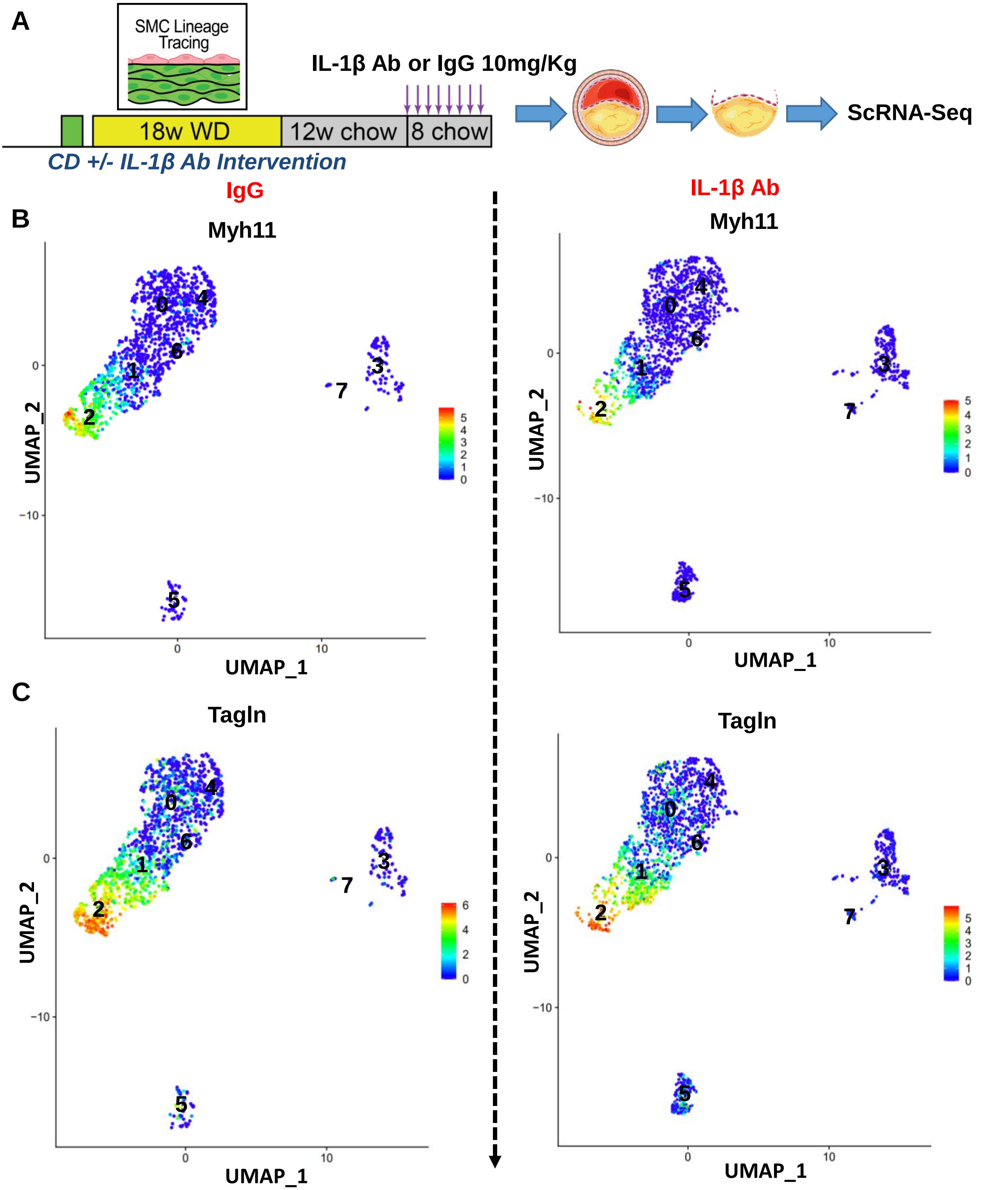


**Supplemental Figure XI. IL-1β Ab Intervention in *Apoe*$^{-/-}$ mice after chow diet-induced lipid lowering resulted in reduced SMC marker+ clusters in the BCA lesions compared to age and diet matched IgG controls. A**, Experimental design, SMC-lineage tracing Apoe−/− mice were injected with tamoxifen at 6 to 8 weeks of age and subsequently placed on a Western diet (WD) for 18 weeks to induce advanced atherosclerosis and randomly distributed into two groups. One group continued receiving WD for another 12 weeks, and another group received chow diet for 12 weeks. Mice were then randomized to being treated with a murine IL-1β neutralizing Ab or an isotype-matched IgG control Ab for 8 weeks while mice continued on chow diet. **B**, Myh11 UMAP and **C**, Tagln UMAP analysis shows cells from IgG on left and from IL-1β on the right, and red arrows indicating number of cells were decreased in cluster 2 of IL-1β Ab group compared to IgG group.

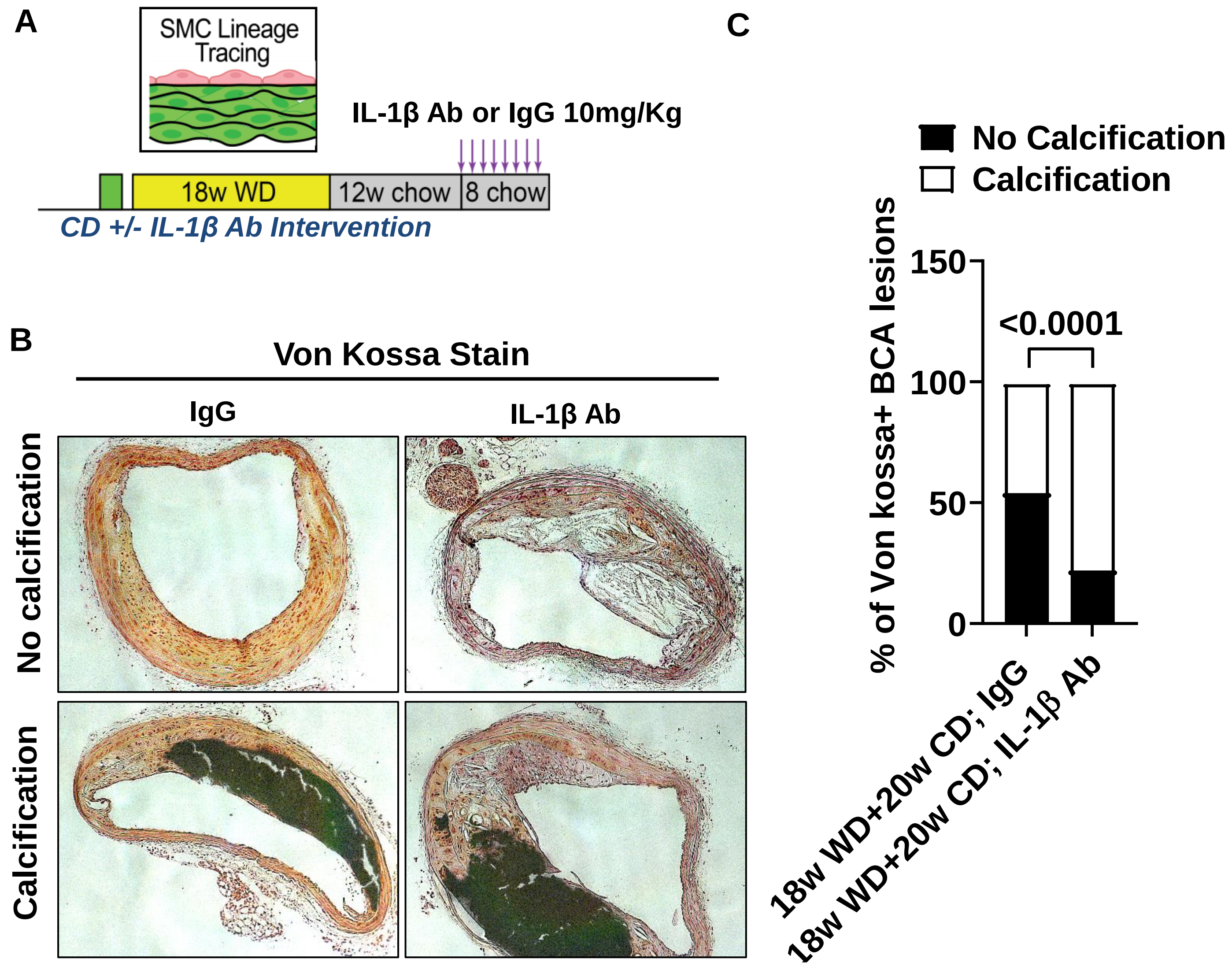


**Supplemental Figure XII. IL-1β Ab Intervention in *Apoe*$^{-/-}$ mice after chow diet-induced lipid lowering did not change calcification in the BCA lesions compared to age and diet matched IgG controls. A**, Experimental design, SMC-lineage tracing Apoe−/− mice were injected with tamoxifen at 6 to 8 weeks of age and subsequently placed on a Western diet (WD) for 18 weeks to induce advanced atherosclerosis and randomly distributed into two groups. One group continued receiving WD for another 12 weeks, and another group received chow diet for 12 weeks. Mice were then randomized to being treated with a murine IL-1β neutralizing Ab or an isotype-matched IgG control Ab for 8 weeks while mice continued on chow diet. **B**, Von Kossa stained representative BCA images. **C**, Percentage of mice showing calcification and no calcification. p-values displayed refer to Fisher's exact test between IgG and IL-1β Ab groups.

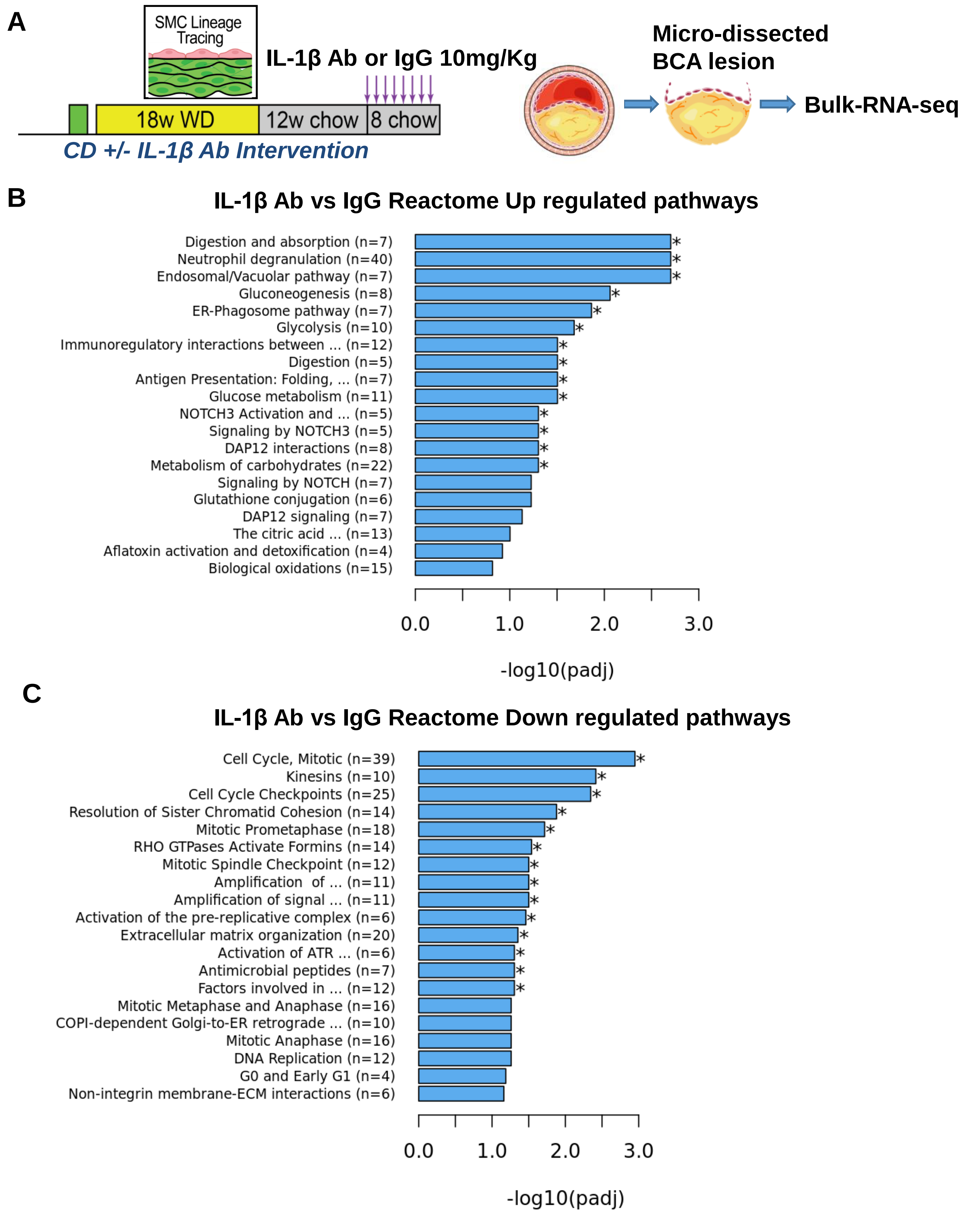


**Figure XIII. Treatment of *Apoe*$^{-/-}$ mice with an IL-1β Ab after chow diet-induced lipid lowering resulted in up-regulated neutrophil degranulation and bioenergetic pathways and down-regulated cell cycle pathways in the BCA lesions. A**, Experimental design, SMC-lineage tracing Apoe−/− mice were injected with tamoxifen at 6 to 8 weeks of age and subsequently placed on a Western diet (WD) for 18 weeks to induce advanced atherosclerosis and randomly distributed into two groups. One group continued receiving WD for another 12 weeks, and another group received chow diet for 12 weeks. Mice were then randomized to being treated with a murine IL-1β neutralizing Ab or an isotype-matched IgG control Ab for 8 weeks while mice continued on chow diet. **B**, Reactome upregulated pathways **C**, Reactome down regulated pathways.

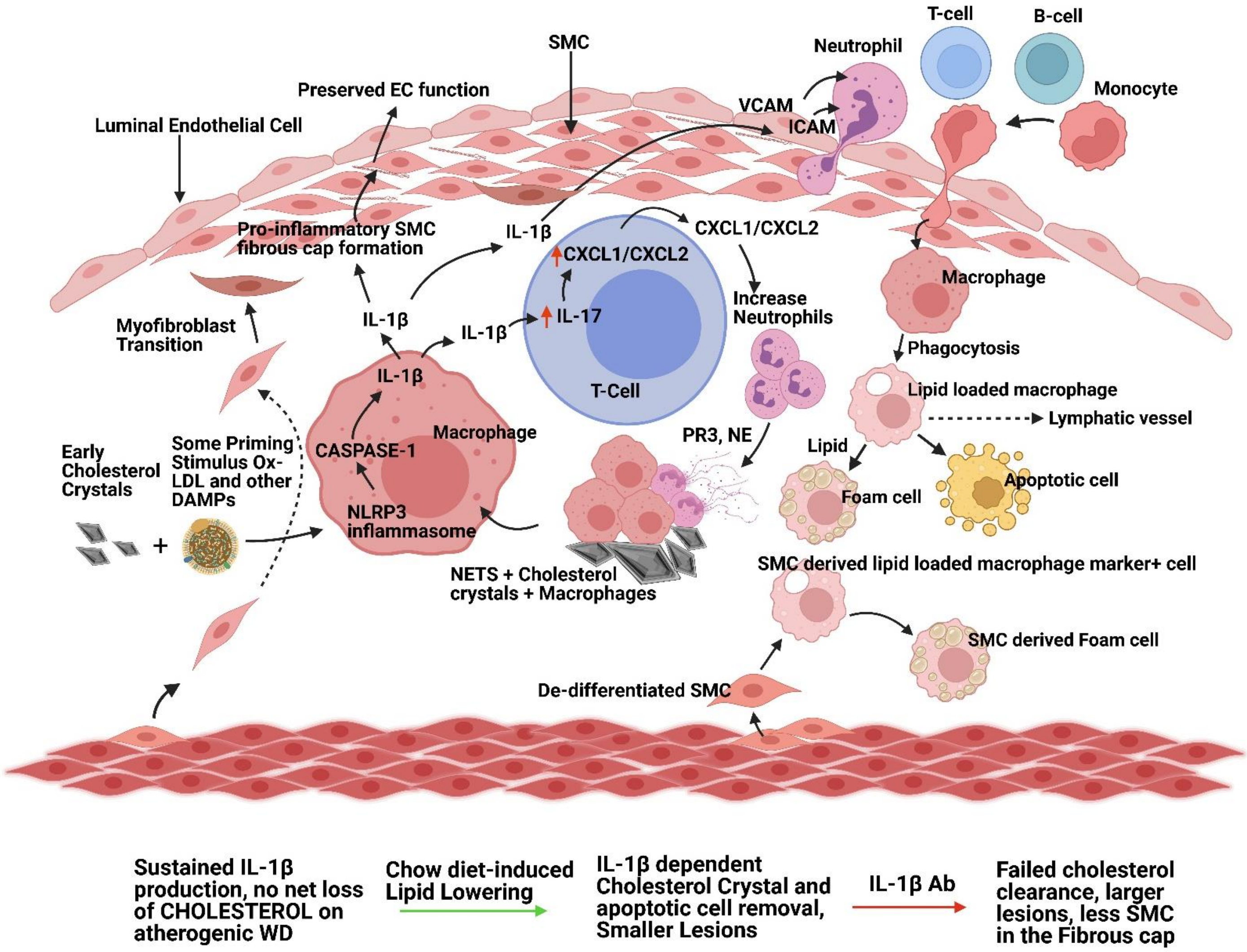


**Figure XIV. Treatment of *Apoe*$^{-/-}$ mice with an IL-1β Ab after chow diet-induced lipid lowering resulted in failed clearance of cholesterol crystals and increased reduced indices of plaque stability**. Evolutionarily conserved IL-1β dependent protective mechanisms act cooperatively to remove cholesterol/cholesterol crystals and apoptotic cells but are over-whelmed and lesions continue to progress as long as mice continue on a WD. Upon switching to chow diet these IL-1β-dependent processes are now sufficient to induce reduced cholesterol content and increased plaque stability. Graphical abstract created with BioRender.com
