## supplemental methods for "IL-1β inhibition partially negates the beneficial effects of diet-induced lipid lowering"

**Supplemental table 1**

| **Metal** | **Marker** | **Clone** | **Vendor** | **Concentration** | **Surface or intracellular** |
|---|---|---|---|---|---|
| Y89 | CD45 | 30-F11 | Fluidigm | 120ng/mL | Surface |
| La139 | CD206 | C068C2 | Biolegend | 600ng/mL | Intracellular |
| Pr141 | CD140b | APB5 | eBioscience | 300ng/mL | Surface |
| Nd142 | TRPV4 | Polyclonal | Invitrogen | 600ng/mL | Intracellular |
| Nd143 | CD117 | 2B8 | Biolegend | 1000ng/mL | Surface |
| Nd145 | Desmin | RD301 | Abcam | 2000ng/mL | Intracellular |
| Nd146 | S100B | 4C4.9 | Abcam | 10000ng/mL | Intracellular |
| Sm147 | CD200 | OX-90 | Biolegend | 300ng/mL | Surface |
| Nd148 | NG2 | 546930 | R&D | 2000ng/mL | Surface |
| Nd150 | Oct3/4 | 40/Oct-3 | BD | 2000ng/mL | Intracellular |
| Sm154 | Runx2 | 232902 | R&D | 1000ng/mL | Intracellular |
| Gd156 | CD184 | L276F12 | Biolegend | 300ng/mL | Surface |
| Gd157 | CD124 | mIL4R-M1 | BD | 30ng/mL | Surface |
| Tb159 | SMA | 1A4 | eBioscience | 4ng/mL | Intracellular |
| Dy161 | CD34 | Ram34 | eBioscience | 250ng/mL | Surface |
| Dy162 | GFP | 338002 | Biolegend | 4000ng/mL | Intracellular |
| Dy163 | CD11b | M1/70 | Biolegend | 250ng/mL | Surface |
| Dy164 | CX3CR1 | SA011F11 | Fluidigm | 0.1uL/200uL | Surface |
| Er167 | IL-6 | MP5-20F3 | Fluidigm | 1uL/200uL | Surface |
| Er168 | CD140a | APA5 | Biolegend | 1000ng/mL | Surface |
| Tm169 | Ly6A/E | D7 | Fluidigm | 0.1uL/200uL | Surface |
| Er170 | IFNg | XMG1.2 | Biolegend | 10000ng/mL | Intracellular |
| Yb171 | CD44 | IM7 | BD | 1000ng/mL | Surface |
| Yb172 | CD86 | GL1 | Fluidigm | 1uL/200uL | Surface |
| Yb174 | Klf4 | Polyclonal | CST | 2000ng/mL | Intracellular |
| Lu175 | F4/80 | BM8 | Biolegend | 360ng/mL | Surface |

**CyTOF data analysis:**

After measurement the samples were normalized[46], de-barcoded[47], and uploaded to Cytobank (www.cytobank.org) for cleanup gating before exporting FCS files for downstream analysis. Raw data was transformed using a division factor of 5[48]. Dimensionality reduction was done by UMAP using default parameters before using the full graph output by UMAP for Leiden clustering with default parameters.
